# An RNA-mediated DNA melting mechanism for CRISPR-Cas9

**DOI:** 10.1101/2025.09.04.674090

**Authors:** George Hedger, Virginia Jiang, Freja Ekman, Qi Wang, David E. Shaw

## Abstract

CRISPR-Cas9 systems, adaptive defense mechanisms in bacteria and archaea, have been widely adopted as powerful gene editing tools, revolutionizing biological and medical research. In the first steps of CRISPR-Cas9 gene editing, the Cas9 protein, in complex with RNA, facilitates DNA melting and subsequent RNA-DNA hybrid formation, but the atomic-level mechanism of this fundamental process is not fully understood. Here, we present the results of long-timescale molecular dynamics simulations in which Cas9-RNA complexes bound to double-helical DNA and promoted the formation of RNA-DNA base pairs in a unidirectional, stepwise manner. Unexpectedly, we observed a direct role for the RNA in facilitating DNA melting events through a mechanism in which RNA bases intercalated within the DNA and promoted strand separation. In addition, breathing motions within the Cas9 DNA-binding cleft contributed to the sequential formation of RNA-DNA base pairs. These simulation results, obtained for two structurally distinct Cas9 proteins, together with supporting experimental work, suggest a novel RNA-dependent mechanism for DNA melting that may be conserved in other Cas proteins.

## Introduction

The clustered regularly interspaced short palindromic repeat (CRISPR)–CRISPR-associated protein 9 (Cas9) system is an adaptive defense mechanism in bacteria and archaea that targets infecting phages and foreign plasmid DNA.^1,2^ The Cas9 protein uses guide RNA to identify and bind to a specific sequence in the invading DNA at which the DNA endonuclease activity of Cas9 generates double-stranded breaks.^3^ The system has been engineered into a powerful and widely used genome editing tool in which a synthetic single-guide RNA (sgRNA) can be designed to target Cas9 to virtually any locus,^4^ at which insertions, deletions, substitutions, or a variety of modifications can be introduced.^5–10^

Cas9 has a bilobed architecture; the recognition (REC) lobe binds sgRNA, and the nuclease (NUC) lobe contains the C-terminal domain (CTD) and two catalytic sites for double-stranded DNA (dsDNA) cleavage (the domain organization is illustrated in Fig. 1A).^1,11,12^ For sgRNA-loaded Cas9 to identify its target sequence, the target must be preceded by a short protospacer adjacent motif (PAM) that is recognized by the PAM-interacting (PI) domain of the Cas9 CTD.^13^ Once the PAM is engaged, the Cas9-sgRNA complex catalyzes strand separation of the DNA and facilitates the formation of an RNA-DNA hybrid in a sequence-specific, unidirectional manner.^13–20^ Strand separation in the middle of B-form DNA involves breaking multiple Watson–Crick base pairs and stacking interactions among the bases—an energetically unfavorable process that creates a kinetic barrier substantially higher than can easily be overcome by thermal fluctuations.^21^ It has been proposed that Cas9 bends the DNA and induces base-flipping in the PAM-proximal region (PAM +1, +2, and +3), and that RNA-DNA base pairing proceeds through the target sequence by a sequential, stepwise unwinding mechanism,^1,13,15,16,20,22–24^ but our atomic-level understanding of this fundamental process is incomplete.

**Fig. 1.**
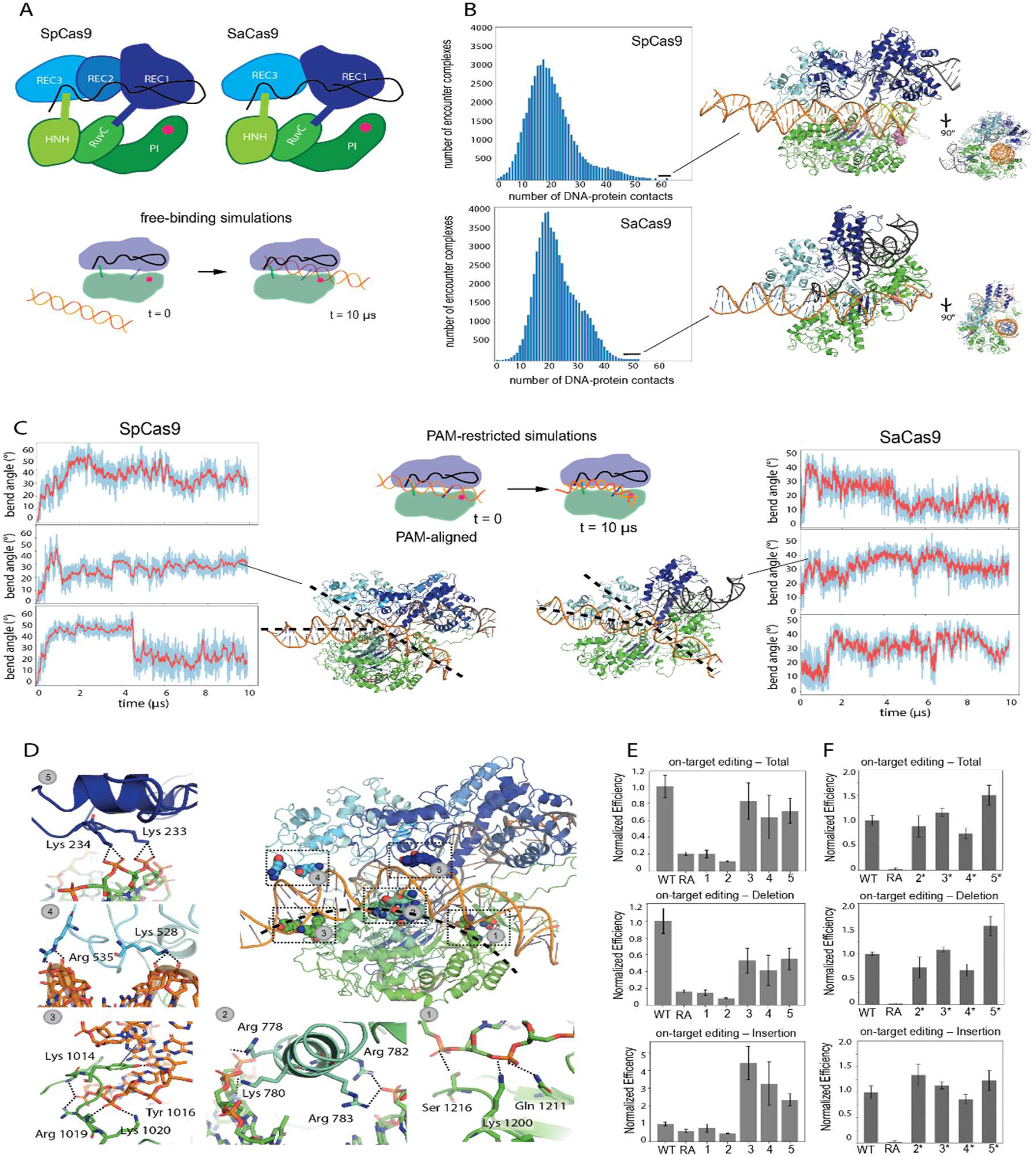
Cas9 binds and bends dsDNA in the groove between the REC and NUC lobes. (A) Top: Cartoon illustration of the domain organization of SpCas9 and SaCas9. Domains in the recognition (REC) lobe and Nuclease (NUC) lobe are colored in blue and green, respectively, the PAM-binding arginines in the PI domain are represented by a pink circle, and the sgRNA is shown as a black line. Bottom: A simplified schematic of the free-binding simulations. (B) Histograms of the number of protein-DNA contacts (by which we mean the number of protein residues in contact with DNA; see methods) observed for sgRNA-Cas9-dsDNA encounter complexes generated from free-binding simulations. On the right are representative structural models of the most stable sgRNA-Cas9-dsDNA encounter complexes (indicated with a bar at the far right of the histograms). (C) The duplex bend angle as a function of time for three replicate simulations of the Cas9 PI domain pre-aligned to the PAM motifs of linear duplexes; SpCas9 on the far left, SaCas9 on the far right. (A simplified schematic of the PAM-aligned simulations is shown at the top middle.) Next to each set of plots is representative structural model of Cas9 stabilizing a bent conformation of the dsDNA. The bend angle was defined as the angle between the phosphorous atoms of tsDNA bases −5, +5, and +25. The raw values are shown in cyan, and the mean of a sliding window of 10 data points is shown in red. (D) Atomic details of how five clusters of residues in SpCas9 (one cluster from each domain that interacts with the DNA) interact with a bent dsDNA. The black dashed line indicates the concave surface of the NUC lobe. (E) Results from a cell-based assay showing on-target editing (total, insertion, and deletion) for WT SpCas9 and five variants, each with one of the clusters noted in panel D mutated: Variant 1 (PI domain): K1200A/S1216A/Q1221A; Variant 2 (HNH domain): R778A/R780A/R783A/K782A; Variant 3 (RuvC domain): K1014A/Y1016A/R1019A/K1020A; Variant 4 (REC3 domain): K528A/K535A/K536A; Variant 5 (REC2 domain): K233A/K234A. RA: R1333A (PAM-binding arginine). Error bars represent standard deviation. (See methods section for details.) (F) Results from a cell-based assay showing total on-target editing (total, insertion, and deletion) for WT SpCas9 and four arginine-substituted variants, each with one of the clusters noted in panel D mutated: Variant 2* (HNH domain): E779R/M781R/K782R; Variant 3* (RuvC domain): K1014R/Y1016R/K1020R; Variant 4* (REC3 domain): K528R/K535R/K536R; Variant 5* (REC2 domain): K233R/K234R/E232R/N235R. Error bars represent standard deviation. (See methods section for details.)

A challenge in characterizing the atomic-level mechanism by which Cas9 binds and deforms dsDNA arises because the transient and dynamic nature of Cas9-DNA encounter complexes makes them difficult to capture using experimental structural approaches. Although most existing structures of Cas9-sgRNA-DNA complexes are of post-hybrid states, and reveal little about what happens prior to or during DNA-RNA hybrid formation, several insightful approaches—such as covalently linking Cas9 to DNA, using DNA with local mismatches, or using less-studied Cas9 variants^22–26^—have provided valuable information about the structural intermediates along the DNA deformation pathway induced by Cas9 from *Streptococcus pyogenes* (SpCas9).^22–24^ Due to the structural diversity of Cas9 family proteins,^27–31^ however, the extent to which the insights gained for SpCas9 apply to other Cas9 family members is unclear. The REC2 domain of the REC lobe, for example, appears to be important for mediating SpCas9-DNA encounter complexes,^22,23^ but *Staphylococcus aureus* Cas9 (SaCas9) lacks a REC2 domain,^29,32^ suggesting a different or modified mechanism might be operative in that context.^22,23,29^

In the work reported here, we used long-timescale molecular dynamics (MD) simulations, along with cell-based and biophysical assays, to study the atomic-level mechanisms of SpCas9^17,33^ and SaCas9^29^—two structurally distinct and widely used Cas9 proteins—starting from their encounter with DNA and proceeding through RNA-DNA heteroduplex formation. In the simulations, Cas9-RNA complexes bound to and melted linear B-form DNA, and subsequently facilitated the formation of RNA-DNA base pairs in a unidirectional, stepwise manner. This result is consistent with what has been observed experimentally; additionally, mutation of certain residues that mediated DNA binding in our simulations impacted editing efficiency in our cell-based assays. The detailed structural mechanism observed in the simulations was, however, unexpected. Notably, sgRNA entered into Mg^2+^-mediated interactions with the DNA, allowing sgRNA bases to intercalate into the DNA and induce local and reversible DNA base-pair breaks.

This finding is consistent with our biophysical experiments, which show that single-stranded RNA (ssRNA) can impact the melting profile of the dsDNA. To our knowledge, no previous study has suggested a role for Cas-bound RNA in DNA deformation, but our results provide evidence that the sgRNA has a direct role in DNA melting in SaCas9 and SpCas9, a mechanism that may be shared by other Cas family members. The simulations also suggest that breathing motions of the groove formed by the REC and NUC lobes fine-tune the distance between the bases of the sgRNA and the partially melted target strand of DNA (tsDNA). This promotes in-register RNA-DNA base pairing starting from the first PAM-proximal base of the target sequence, and breaks base pairs that form prematurely further downstream—a potential proofreading mechanism to ensure sequential base pairing from the immediate PAM-proximal region.

## Results

### Cas9 binds and bends linear dsDNA in the cleft between the REC lobe and NUC lobes

To study how Cas9 proteins may interact with linear dsDNA, we first performed a series of free-binding simulations starting with a linear B-DNA duplex in solution separated from an sgRNA-loaded Cas9, using PDB ID 4ZT0^20^ for SpCas9 and PDB ID 5CZZ^29^ for SaCas9, with the DNA removed from the latter (see Methods for details of all simulations). For each Cas9 protein, we performed 20 independent 5 μs simulations and observed the spontaneous formation of a variety of Cas9-DNA encounter complexes (Fig. S1A). We used the number of protein residues contacting the DNA as a metric to judge the stability of these encounter complexes (Fig. 1B, left), and representative structural models of the most stable sgRNA-Cas9-dsDNA encounter complexes are shown on the right side of Fig. 1B.

For both Cas9 proteins, we observed that in the most stable DNA encounter complexes the DNA was bound in the cleft between the REC and NUC lobes (Fig. 1B; Supplementary Video 1). This DNA binding mode is reminiscent of what is observed in post-hybrid Cas9 crystal structures^11,17^ (Fig. S1B). We refer to this Cas9 conformation as “open-Cas9” because the distance between the REC3 domain of the REC lobe and the RuvC domain of the NUC lobe is larger than that in the states prior to DNA binding (“closed-Cas9”), and thus can accommodate binding of dsDNA or a DNA/RNA hybrid. Both the open and closed conformations of Cas9 were repeatedly visited in the simulations (Fig. S1B), and the dsDNA bound to both of these conformations. Transitions between the open and closed states were not observed while the dsDNA was bound, and shifts in the distribution between the open and closed states in the presence of dsDNA likely resulted from incomplete convergence of the simulations (Fig. S1B). In none of these encounter complexes, however, was the PAM sequence (NGG for SpCas9 and NNGRRT for SaCas9) correctly aligned with the two arginine residues on the PI domain known to be important for Cas9 recognition of DNA.^17,29^

To capture more biologically relevant encounter complexes in our simulations, we applied a distance restraint to maintain the PAM sequences in close proximity (<5 Å) to the corresponding arginine residues in the PI domain (at the beginning of the simulations, the rest of the DNA was not interacting with any part of the protein) (Fig. 1C). For both Cas9 proteins, as we had observed in the free-binding simulations, the most stable DNA encounter complexes had the DNA bound in the cleft between the REC lobe and the NUC lobe in the open-Cas9 conformation. The bound DNA exhibited spontaneous and reversible bending on the order of tens of microseconds (Fig. 1C; Supplementary Video 2); the angular deviation from linear reached 20°–50° (Fig. 1C). The bending occurred within the interlobe groove of Cas9, placing the PAM-distal DNA base pairs (PAM +4 through +20) between the RuvC domain and the REC3 domain (Fig. 1D and Fig. S1C). Because spontaneous DNA binding and bending was observed for both Cas9 proteins, but with different residue-DNA interactions for each, it is likely that this process derives from the domain-scale architecture of the complexes rather than from specific interactions.

The observed bending was accompanied by the formation of a number of DNA-protein contacts that are not possible for linear DNA to form. These contacts were able to occur because the convex surface of the NUC lobe interacted extensively with the DNA surface and served as a scaffold to stabilize the curved dsDNA (Fig. 1D and Fig. S1C). In addition to the encounter complexes formed with open-Cas9, the dsDNA formed a variety of other complexes with Cas9 in less open conformations, including one resembling the closed Cas9 conformation observed in the PDB ID 4ZT0 crystal structure (Fig. S1, A and D), but fewer DNA-protein contacts formed in these complexes (44 at most) than in the open-Cas9/dsDNA complexes (up to 65) (Fig. S1D). For this reason, we chose to characterize only the open-Cas9/dsDNA complexes in this paper. We note, however, that the alternative Cas9-dsDNA complexes may also be relevant, as they might represent intermediate structures along the dsDNA deformation pathways (see Discussion).

### Mutation of residues that mediated the bent DNA-Cas9 encounter complexes in simulations altered SpCas9 editing efficiency in cells

We next probed the significance of different open-Cas9/dsDNA interactions using an assay in which we measured the activity of numerous SpCas9 variants in human embryonic kidney cells. The on-target editing efficiency, insertion efficiency, and deletion efficiency of SpCas9 were measured through next-generation sequencing, and the activity levels were normalized against protein expression levels, as quantified by western blot (Fig. S1E). Wild-type (WT) SpCas9 had an overall on-target editing efficiency of 72% ± 11%, averaged over three independent experiments. More than 85% of the on-target editing events resulted in partial deletion in the targeted gene, and in less than 10% of events a non-templated small-gene insertion in the targeted region was also detected. As expected, mutation of one of the PAM-coordinating arginines in the PI domain (R1333A) substantially decreased SpCas9 activity, giving an overall on-target editing efficiency of 15% ± 7% (Fig. 1E).

We next used this assay to evaluate the functional consequences of independently mutating clusters of residues that mediated protein-DNA contacts in our simulations (one cluster from each SpCas9 structural domain that made contact with the DNA), and found that the impact on on-target editing efficiency varied depending on the mutated cluster. The variant with mutations in the HNH domain (R778A/R780A/R783A/K782A) had very low on-target editing efficiency, similar to what was observed for the PAM-binding defective mutant R1333A (Fig. 1E), despite the fact that the expression level of this mutant was comparable to that of the WT control. This residue cluster is located on a short helix at the N-terminus of the HNH domain and is distal to the HNH domain catalytic site, so the loss-of-function is unlikely to be explained by direct disruption of HNH catalytic activity. In addition, the PI domain variant (K1200A/S1216A/Q1221A), which includes mutation of the functionally important S1216 residue,^34,35^ also resulted in on-target editing efficiency on par with the PAM-binding defective mutant.

Variants with mutations in the REC2 domain (K233A/K234A), REC3 domain (K528A/K535A/K536A), and RuvC domain (K1014A/Y1016A/R1019A/K1020A) had more moderate effects, with each variant causing a 20%–40% reduction in the overall on-target editing efficiency. This decrease in editing efficiency was primarily associated with a decrease in the deletion activity (Fig. 1E). Interestingly, in both the RuvC and REC3 domain variants, we observed over a 3-fold increase in insertion editing efficiency relative to the WT protein. We have not studied the mechanisms that underlie differential control of Cas9 insertion and deletion activity, but note that the decrease in deletion activity cannot be explained by lower expression or improper folding of the mutant proteins, common mechanisms that often account for decreased activity in mutants.

In addition to testing the impact of alanine mutations designed to weaken dsDNA-Cas9 interactions, we also tested arginine mutations in clusters of residues in four of these five domains that we predicted might enhance dsDNA-Cas9 interactions. (We did not assess mutations in the PI domain cluster because it is adjacent to the PAM-binding residues, where arginine mutations could interfere with PAM recognition.) Variants with arginine mutations in the HNH domain (E779R/M781R/K782R), REC3 domain (K528R/K535R/K536R), and RuvC domain (K1014R/Y1016R/ K1020R) showed no significant differences in editing efficiency from the WT protein. In contrast, arginine mutations in the REC2 domain (K233R/K234R/E232R/N235R) led to a ∼50% increase in normalized overall on-target editing efficiency, primarily driven by an increase in deletion efficiency (Fig. 1F and Fig. S1F).

Because gain-of-function mutations are inherently more challenging to design than loss-of-function mutations, further efforts, such as computationally guided sequence optimization or saturation mutagenesis, would likely be necessary to further enhance activity.

### In simulations, sgRNA interacts with dsDNA through Mg^2+^-mediated interactions and promotes transient dsDNA base-pair breaks

We next performed 20 50-µs simulations initiated from dsDNA-Cas9 encounter complexes generated by our previous simulations in which the PAM had been restrained (10 simulations each for SpCas9 and SaCas9). In addition to again observing the protein-dsDNA interactions discussed above, we also observed Mg^2+^-mediated sgRNA interactions with the dsDNA in all 20 of these simulations (Fig. 2A and Fig. S2A). These interactions relied on the presence of Mg^2+^, which chelated the backbone phosphates of both molecules; removing Mg^2+^ from the simulations in regions where both backbones were chelated immediately triggered separation of the sgRNA from the dsDNA. The number of Mg^2+^ ions bound between the dsDNA and sgRNA ranged from one to eight in individual simulations (Fig. S2B), and binding occurred downstream of the PAM-proximal region. Once an Mg^2+^ ion became bound at the sgRNA-dsDNA interface, it remained stably bound until the end of the simulation. Although Mg^2+^ has been resolved in some sgRNA-Cas9-DNA X-ray structures (e.g., PDB ID 6O0X^11^), Mg^2+^-mediated dsDNA-sgRNA interactions have not been reported previously for Cas9.

**Fig. 2.**
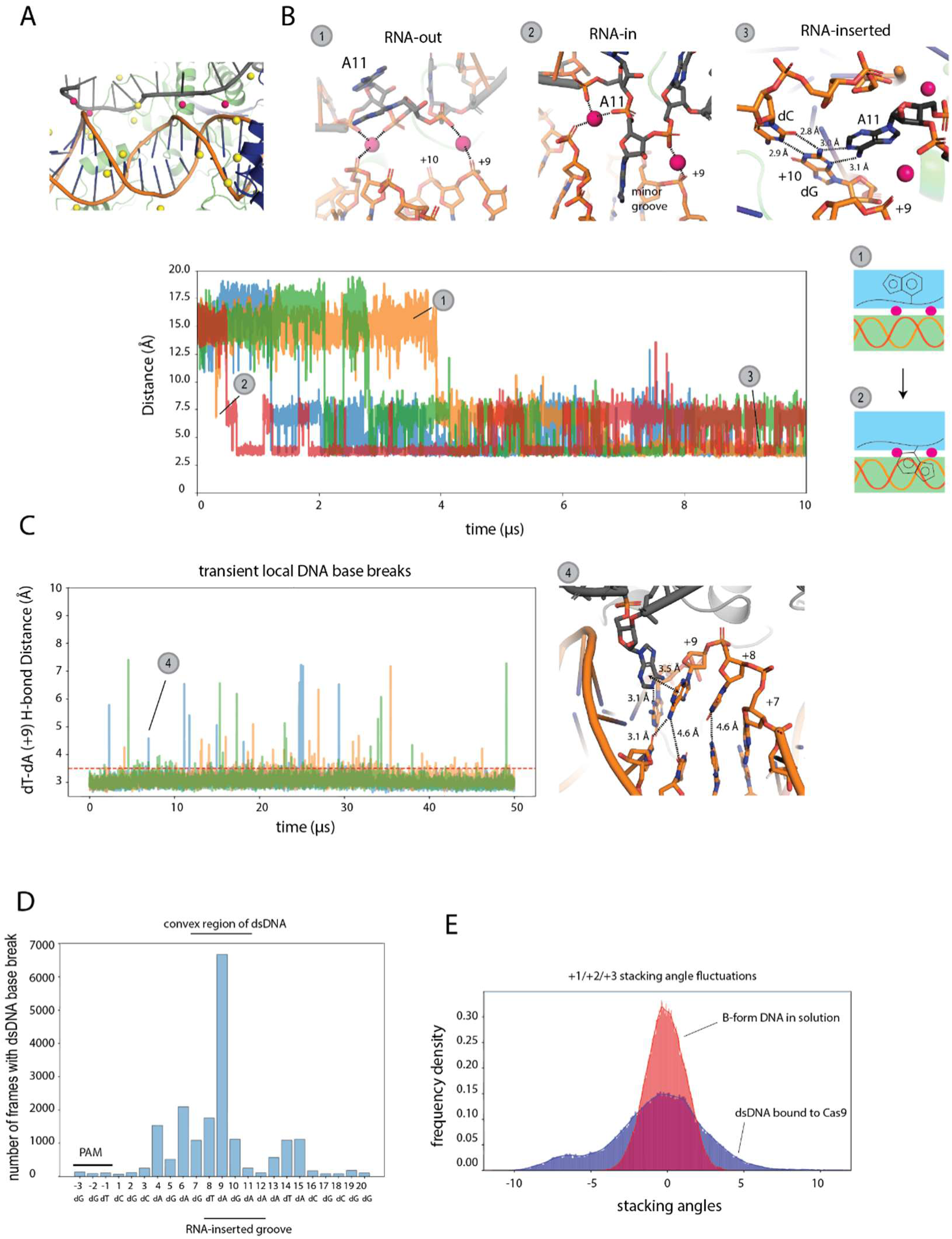
Mg^2+^-mediated sgRNA-dsDNA interactions in SpCas9-dsDNA encounter complexes. (A) A representative structure of Mg^2+^ ions binding between the sgRNA (grey) and the dsDNA (orange). Pink spheres are Mg^2+^ ions mediating backbone interactions; yellow spheres are non-interaction-mediating Mg^2+^ ions. (B) Top: Atomic details of Mg^2+^-mediated sgRNA-dsDNA backbone interactions, with Mg^2+^ ions represented as pink spheres. In state 1 (RNA-out), the RNA has flipped to the RNA-out state; in state 2 (RNA-in), the flipped RNA base has entered the minor groove of the DNA without forming hydrogen bond interactions; and in state 3 (RNA-inserted), the RNA base has hydrogen bonded with a dsDNA base in the minor groove. These structures are derived from instantaneous frames from one of the simulations used for the plot in the bottom portion of this panel. Bottom left: Time course showing the hydrogen-bonding distance between A11 of the sgRNA and dG(+10) of the tsDNA over the course of four replicate Cas9-dsDNA encounter complex simulations, each colored differently, in which the RNA was observed to interact with the minor groove of the dsDNA. Bottom right: Cartoon illustration of the conformational transition from the RNA-out to the RNA-inserted state. Mg^2+^ ions are represented as pink circles. (C) Time course of the distance between the +9 DNA bases for four replicate RNA-inserted Cas9-dsDNA encounter complex simulations, each colored differently; transient base-pair breaks at +9 were observed. A representative structure of RNA stabilizing a transiently broken base (state 4) is shown on the right and is identified in the time course. (D) Distribution of base-pair break events along the Cas9-bound DNA region in RNA-inserted Cas9-dsDNA encounter complex simulations. (E) Distribution of the dsDNA stacking angles at the PAM-proximal region (PAM +1, +2, and +3) in RNA-inserted Cas9-dsDNA encounter complex simulations (blue) and for dsDNA alone in solution (red).

In five of the 20 simulations (four for SpCas9 and one for SaCas9), we observed individual sgRNA nucleotides flip from their original REC-domain-facing positions, the “out” position, to an “in” position from which they could intercalate into the minor groove of the dsDNA (Fig. 2B and Fig. S2C; Supplementary Video 3). Such intercalation of an sgRNA base into the minor grove was observed at multiple locations on the dsDNA. In the intercalated state, the flipped RNA base formed non–Watson-Crick hydrogen bonds with DNA bases and disturbed stacking with adjacent bases (Fig. 2B and Fig. S2C). In simulations in which sgRNA base A11 was observed to be intercalated at the PAM +9 position of the dsDNA, we also observed transiently stable structures in which the complementary DNA bases at PAM +9 were separated by more than 4.5 Å (Fig. 2C). In such cases, the PAM +9 base of the tsDNA formed a transiently stable non–Watson-Crick hydrogen bond with the intercalated RNA base (Fig. 2C). This suggests that the sgRNA can stabilize partially broken dsDNA structures. Furthermore, although the entire dsDNA remained highly helical during the aggregate ∼1.5 ms of simulation time of these 20 simulations, we found that such transient dsDNA base-pair breaks happened not only at the regions of RNA intercalation, but also throughout the entirety of the dsDNA bound within Cas9, including in the PAM-proximal region (Fig. 2D and Fig. S2D).

In addition to the observed base-pair breaks, the stacking angles of the bases in the PAM-proximal region also frequently deviated from the ideal near planar (<6°) that is observed for dsDNA alone (Fig. 2E and Fig. S2E). The PAM +9 position also happened to be at the curved region of the bent dsDNA (bases +9 to +13) that resulted from Cas9 binding. The frequency of transient base-pair breaks depended on both the severity of the DNA bending and on the intercalation of the RNA base (Fig. S2F), suggesting that the observed transient local base-pair breaks likely resulted from a composite effect of both phenomena. In addition to the minor-groove intercalation, we also observed individual sgRNA bases interact with the backbone phosphates of the dsDNA directly or through Mg^2+^-mediated interactions (Fig. S2G). Such interactions may also impact dsDNA stability.

### An ssRNA promotes dsDNA separation in simulations and alters the melting profile of dsDNA in biophysical experiments, and these effects can be further sensitized by Mg^2+^

The simulations above suggest that sgRNA not only serves as the sequence-specific guide for Cas9 to identify its target, but that prior to heteroduplex formation it may also promote reversible dsDNA base-pair breaks at the RNA-binding site as well as reversible dsDNA strand separation distal to the RNA-binding site. To provide further evidence for a role of Mg^2+^ and sgRNA in promoting dsDNA deformation, we designed a mini sgRNA-Mg^2+^-dsDNA complex consisting of only a five-base strand of sgRNA (which we will refer to as “ssRNA-5”; sequence of AAAGA, complementary to DNA bases +9 through +13) intercalated into a twenty-base-pair segment of dsDNA, and simulated this complex in the absence of Cas9 (Fig. 3A). In 18 of 20 such simulations, the dsDNA base pairs adjacent to the intercalated RNA base separated, resembling the transiently broken structures from our simulations of sgRNA-loaded Cas9 bound to dsDNA (Fig. 3A and Fig. S3A). Unexpectedly, this local melting was followed by a dramatic shift in the angle of DNA bending, which led to base-pair breaks not only adjacent to the RNA-inserted position, but also extending to the ends of the linear DNA, eventually leading to a more extensive strand separation of the 20-bp DNA fragment (Supplementary Video 4). In the absence of ssRNA-5, however, over an aggregated 400 µs simulation time, we only observed the expected reversible base-pair breaks at the termini of the dsDNA (Fig. 3A and Fig. S3B). These simulation results strongly suggest that the Mg^2+^ and ssRNA can actively promote melting of dsDNA in the absence of Cas9.

**Fig. 3.**
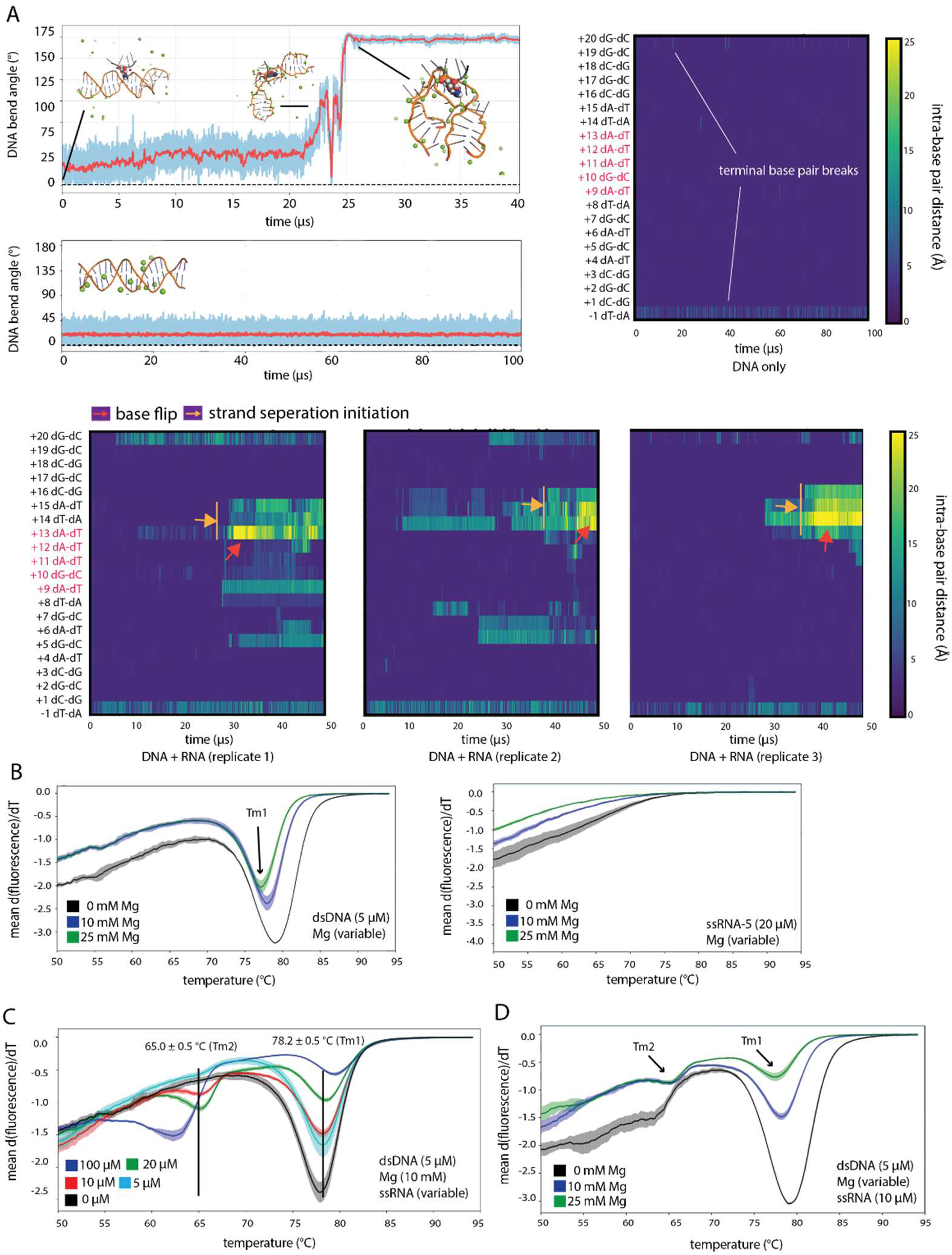
ssRNA promotes dsDNA melting. (A) Top left: Upper plot shows DNA bending in a simulation of an Mg^2+^-chelated ssRNA-dsDNA complex and lower plot shows the bend angle of dsDNA simulated on its own in aqueous solution. Representative structural models from the simulations are shown as insets. Bottom: Base-pair distance analyses of three replicate simulations of ssRNA-dsDNA complexes. The red arrows indicate that the DNA base has flipped out and is exposed for RNA paring. The orange arrows indicate the initiation of strand separation, in which multiple adjacent DNA base pairs are broken. Upper right: Base-pair distance analysis of one replicate simulation of dsDNA alone. The base pairs interacting with the RNA throughout the simulation are labelled in red. (B) Thermostability analysis of a dsDNA (left) and an ssRNA (right) in the presence of variable Mg^2+^ concentrations. (C) Thermostability analysis of the dsDNA in the presence of 10 mM Mg^2+^ and variable ssRNA concentrations. (D) Thermostability analysis of the dsDNA in the presence of 10 mM ssRNA and variable Mg^2+^ concentrations. Standard deviations for the thermostability assays, which were performed in duplicate, are indicated by the lighter-colored bars around the mean.

Motivated by this result, we experimentally measured the thermostability of a 21-bp dsDNA in the presence and absence of both a 21-base ssRNA (which we will refer to as “ssRNA-21”) and Mg^2+^ to directly assess the significance of ssRNA and Mg^2+^ interactions on dsDNA stability. In the absence of Mg^2+^, when the temperature ramped from 27 °C to 95 °C, the melting curve of the dsDNA alone (at 5 µM) featured a single peak, with the T_m_ determined as 78.9 ± 0.5 °C (Fig. 3B). As expected from a previous study,^36^ adding physiologically relevant concentrations of Mg^2+^ to the buffer decreased the T_m_ slightly; at 10 mM Mg^2+^, the T_m_ decreased by about 1.1 °C to 77.8 ± 0.5 °C, and was further decreased to 76.9 ± 0.5 °C at 25 mM Mg^2+^ (Fig. 3B). There were no visible decreases in the derivative of the mean fluorescence over the tested temperature range when assessing ssRNA-21 (without dsDNA) in the presence (10 mM or 25 mM) or absence of Mg^2+^ (Fig. 3B).

We next measured melting curves of the dsDNA in the presence of both ssRNA-21 and Mg^2+^, both at various concentrations, and found that the presence of the ssRNA at higher concentrations drastically changed the melting profile of the dsDNA, consistent with predictions from our simulations. Specifically, although adding 10 mM Mg^2+^ and 5 µM ssRNA-21 (a 1:1 ratio of ssRNA:dsDNA) to the buffer had a minimal effect on the melting curve, increasing the ssRNA concentration to 10 µM (a 2:1 ratio of ssRNA:dsDNA) led to two peaks in the melting curve, suggesting there were two distinct melting events in the sample (Fig. 3C and Fig. S3C).

The major peak was at 78.2 ± 0.5 °C (T_m_1), which is consistent with the peak in the dsDNA-Mg^2+^ melting profile; an additional peak was also observed, however, at 65.0 ± 0.5 °C (T_m_2), nearly 13 °C lower than the major peak. Further increasing the ssRNA concentration led to greatly intensified signal for the second peak, coupled with a decreased signal of the first peak. Strikingly, at 100 µM ssRNA (a 20:1 ratio of ssRNA:dsDNA), the second, lower-temperature peak became the major peak in the melting curve. When the Mg^2+^ concentration was increased to 25 mM, we observed a similar two-transition melting profile, as seen with 10 mM Mg^2+^ (Fig. 3D). T_m_2 appears to also depend on the ssRNA sequence being complementary to the target DNA sequence, as the lower-temperature peak diminished—across a range of Mg²⁺ concentrations—when a sequence-scrambled ssRNA was used (Fig. S3E).

Completely removing Mg^2+^ from the buffer decreased the effect of ssRNA-21 on the dsDNA melting profile; at 10 µM ssRNA in the absence of Mg^2+^, we no longer observed a second peak in the T_m_ curve (Fig. 3D). Although the ssRNA was still able to induce the lower temperature peak at higher (20 µM and 100 µM) RNA concentrations, the higher temperature peak in these melting curves remained the larger temperature peak (Fig. S3D). These results suggest that Mg^2+^ is not essential for the observed ssRNA-mediated dsDNA melting behavior, but sensitizes dsDNA’s response to ssRNA—a finding that has not been reported previously, and is consistent with our observations in simulations.

RNA-DNA heteroduplexes are generally more thermodynamically stable than dsDNA,^37^ so it is expected that an ssRNA can destabilize complementary dsDNA through a mechanism in which initial melting of the DNA occurs spontaneously rather than through a process mediated by the RNA. This mechanism alone, however, cannot fully explain our unexpected observation that RNA-dependent dsDNA melting is sensitive to the presence of Mg²⁺. Rather, our simulations suggest that this phenomenon is best explained by an Mg²⁺-mediated, RNA-driven DNA melting mechanism.

Our thermostability assays probe an equilibrium effect and do not directly address Mg²⁺’s role in regulating the kinetics of the DNA melting observed in our simulations. Prior experimental studies have demonstrated, however, that Mg²⁺ plays a crucial role in modulating the kinetics of DNA binding and R-loop formation within sgRNA-loaded Cas9 complexes. In one case, stopped-flow experiments measuring the kinetics of dsDNA binding to sgRNA-loaded SpCas9 in the presence and absence of Mg²⁺ revealed a nearly 700-fold increase in the association constant when Mg²⁺ was present, whereas the overall stability of the ternary complex remained unchanged in the presence of Mg²⁺.^38^ In a separate study, fluorescent nucleotide–based unwinding experiments measuring R-loop formation kinetics of GeoCas9 (a Cas9 variant from *Geobacillus stearothermophilus*) at different Mg²⁺ concentrations showed that R-loop formation was significantly slower at lower concentrations of Mg²⁺, demonstrating that dsDNA unwinding strongly depends on high Mg²⁺ levels.^39^ Although our simulations cannot definitively determine the molecular mechanism underlying Mg²⁺’s profound effect on DNA binding kinetics and R-loop formation, the consistency of these previously reported kinetic studies with our models strengthens our confidence in the simulation-based observations.

### In simulations, unidirectional, sequential DNA-RNA hybrid formation initiates from partially melted dsDNA structures at the PAM +1 position

Our finding that an ssRNA can substantially change the melting profile of a dsDNA in the presence of Mg^2+^ leads to the question of whether partially melted dsDNA structures, with multiple base-pair breaks, exist transiently while bound with Cas9. Although such partially melted dsDNA structures have not been observed experimentally, transient states are often difficult to capture using experimental structural biology methods. Below, we present a set of simulations suggesting that partially melted dsDNA structures can exist while bound with Cas9, and further that these complexes can progress to unidirectional, sequential RNA-DNA base pairing starting from the PAM +1 position, consistent with conclusions drawn from experimental single-molecule studies.^13,15,16^

The simulations presented above revealed only local transient base-pair breaks when dsDNA was bound to Cas9, and we did not observe extended strand separation within the aggregate millisecond of simulation time. We thus applied an enhanced sampling protocol (see Methods for details) that allowed for more efficient sampling of the reversible dsDNA breaks. This technique tempered (intermittently weakened) the charged interactions between the two strands in the dsDNA, thereby increasing the frequency of DNA base-pair breaks and accelerating the exploration of different DNA-DNA interactions. The interactions between DNA and RNA, and between nucleotides and proteins, were not tempered, so remained the same strength throughout the simulations.

Using this enhanced sampling protocol, we initiated 30 simulations (20 for SpCas9 and 10 for SaCas9) from bent, Cas9-bound dsDNA structures extracted from frames in our earlier simulations (without tempering). In the tempering simulations, we observed various partially and reversibly melted dsDNA structures (Fig. S4, A and B) characterized by extensive strand separation in which the complementary bases of at least three sequential base pairs were at least 5 Å apart. These structures maintained a near helical shape, which we refer to as the “molten-helical” state of the DNA. We also identified six simulations (two for SpCas9 and four for SaCas9) in which at least three in-register tsDNA-sgRNA base pairs were formed starting at PAM +1 following partial melting of the dsDNA (see Fig. 4, A and B, Fig. S4C, and Supplementary Videos 5 and 6). Remarkably, in all of these six simulations, the tsDNA-sgRNA base pair formation proceeded sequentially to +2 and +3, with pairing all the way up to +4 in one of the simulations. The tsDNA-sgRNA structures formed at the end of these six simulations overlay near perfectly with the respective post-heteroduplex formation X-ray structures for SpCas9 (PDB ID 5F9R^17^) and SaCas9 (PDB ID 5CZZ^29^), with an average RMSD between the simulated and observed structures smaller than 2.5 Å (Fig. 4, A and B; Supplementary Videos 5 and 6).

**Fig. 4.**
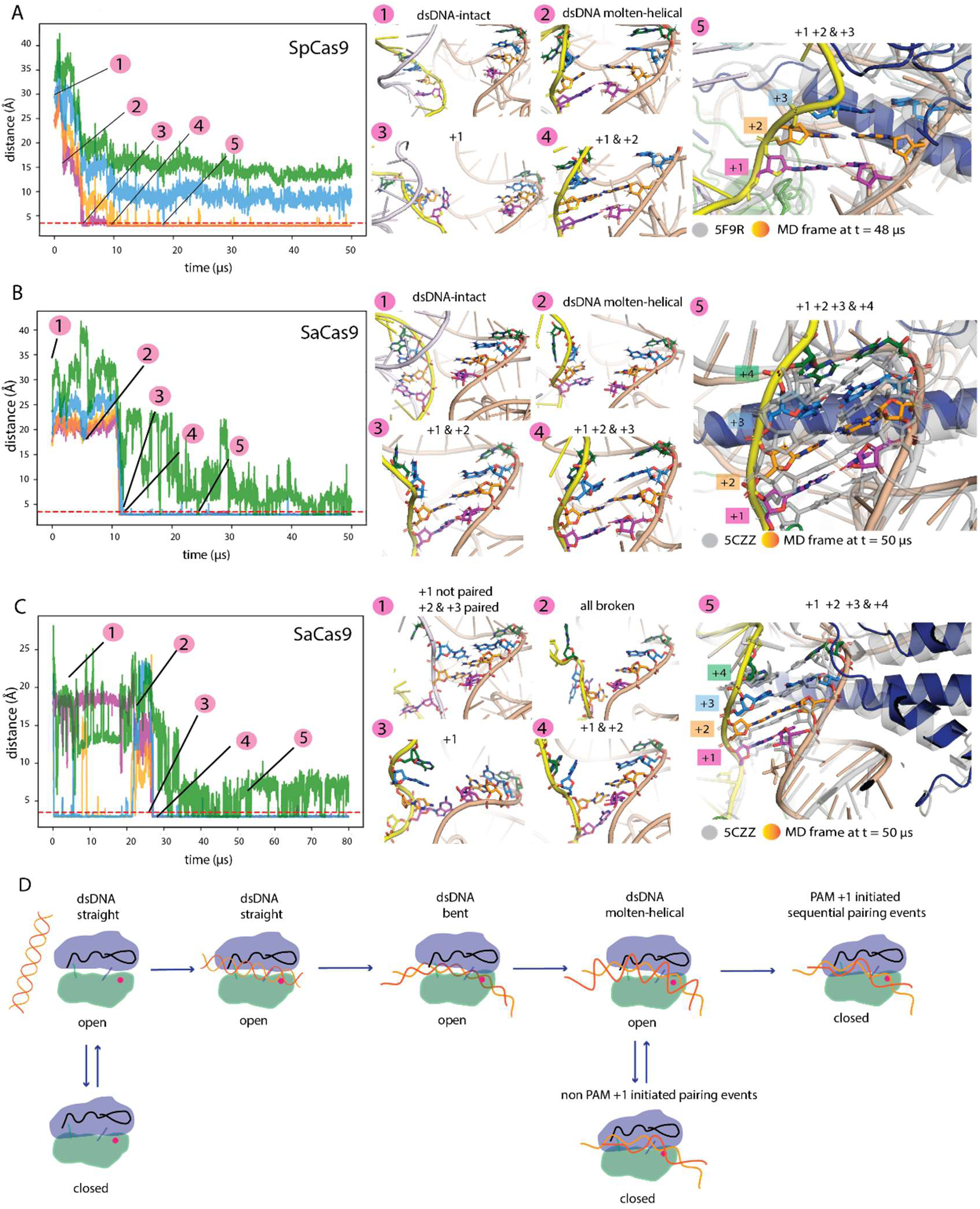
PAM +1-initiated, stepwise heteroduplex formation in simulations of sgRNA-Cas9-dsDNA encounter complexes. (A) A simulation of an sgRNA-SpCas9-dsDNA complex showing sequential, in-register base pairing observed from the +1 position to the +3 position. The distance between the tsDNA base and the corresponding sgRNA base are plotted; +1 shown in magenta, +2 in orange, +3 in blue, and +4 in green. The dashed red line is plotted at a distance of 3.5 Å. The atomic details of structural intermediates occurring in the simulations are shown just to the right of the plot. The +1/+2/+3-paired MD-derived structure is shown overlaid with the crystal structure (PDB ID 5F9R; grey) at the far right. (B) A simulation of an sgRNA-SaCas9-dsDNA complex showing sequential, in-register base pairing observed from the +1 position to the +4 position. The atomic details of structural intermediates occurring in the simulations are shown just to the right of the plot; bases are colored as in panel B. The MD-derived structure paired from +1 to +4 is shown overlaid with the crystal structure (PDB ID 5CZZ) at the far right. (C) A simulation of an SaCas9-dsDNA complex showing the hybrid structure first form base pairs out of sequence (i.e., at +2 rather than +1), but the complex breaks the out-of-sequence base pairs, and subsequently forms sequential base pairs from +1 to +4. The atomic details of structural intermediates occurring in the simulations are shown just to the right the plot; bases are colored as in panel B. The MD-derived structure paired from +1 to +4 is shown overlaid with the crystal structure (PDB ID 5CZZ) on the far right. (D) Cartoon illustration of Cas9-mediated DNA deformation and heteroduplex formation. Cas9-sgRNA first binds and bends dsDNA in the interlobe groove, facilitating local dsDNA melting through an RNA-mediated mechanism. This induces a molten-helical conformation of the DNA from which interdomain motions within Cas9 disfavor DNA-RNA base pairs that are not initiated at PAM+1, and promote PAM +1–initiated, sequential DNA-RNA base pairs. Domains in the recognition (REC) lobe and Nuclease (NUC) lobe are colored in blue and green, respectively, the PAM-binding arginines in the PI domain are represented by a pink circle, and the sgRNA is shown as a black line.

### The closed-to-open conformational transition of Cas9 can break RNA-DNA hybrid structures initiated downstream of PAM +1, an apparent proofreading mechanism to ensure initiation of hybrid formation at PAM +1

In our simulations, each hybrid formation event initiated from PAM +1 involved both flipping of the tsDNA base toward the sgRNA, and a concomitant decrease in the tsDNA-sgRNA backbone distance. This decrease resulted in part from increased tsDNA flexibility caused by downstream melting and unwinding—which enabled the tsDNA backbone to migrate towards the sgRNA—and by an open-to-closed conformational transition of the REC and NUC lobes that brought the sgRNA closer to the tsDNA (Fig. S4E). The RNA-DNA hybrid formation resembles a zipper closing without the slider; the formation of one RNA-DNA base pair decreases the distance between the next two complementary nucleotides and stabilizes the nascent pair by providing a stacking interaction on top of it. In the partially melted dsDNA, unstacking and unwinding was observed along the double-helical structure such that many tsDNA bases were accessible for potential hybrid base pair formation. Indeed, we observed in some simulations that base pair formation occurred first in the PAM-distal region of the DNA (in these cases, the first hybrid base pairs usually formed around PAM +11), but these base pairs were usually only transiently stable and could not serve as a nucleation point for further RNA-DNA hybridization (Fig. S4D).

The PAM +1 position was the most common nucleation site for heteroduplex formation in our simulations. Once the +1 position was engaged, we observed RNA-DNA base pair formation up to the +3 position with SpCas9 and to the +4 position with SaCas9. In one simulation with SaCas9, we observed a hybrid structure first initiate from +2, and for the first 20 µs of the simulation only +2 and +3, but not +1, were paired (Fig. 4C). This simulation gave us an opportunity to observe what might happen when base pairing does not initiate strictly at +1.

Although the +1 disengaged structure remained stable for more than 20 µs, the heteroduplex did not propagate further downstream (i.e., no base pairing was observed at +4 or beyond), and the base pairs at the +2 and +3 position eventually broke at 30 µs. Shortly after the base-pair breaks, the +1 position was able to form an in-register Watson–Crick base pair, followed by pairing and stacking at +2, +3, and +4 (Fig. 4C; Supplementary Video 7). Interestingly, the base-pair breaks at +2 and +3 happened at a time that the REC and NUC lobes were moving from a closed conformation to a more open conformation, physically pulling away the already paired +2 and +3 base pairs and potentially contributing to the base-pair breaks (Fig. S4E). Although such closed-to-open conformational transitions of Cas9 happened in all of our simulations, we never observed breaking of the PAM +1 base pair once it formed, suggesting that the +1 hybrid base pair, compared to those in other positions, is less prone to break as a result of the fluctuation caused by the conformational changes of Cas9. This result suggests a structure-based proofreading mechanism to ensure PAM +1–initiated hybrid formation. We believe it is likely that if the system is able to correct in-register but out-of-sequence Watson–Crick base pairs, it can also correct in-register, mismatched base pairs.

## Discussion

In this study, we used MD simulations, cell-based editing assays, and thermal shift assays to probe the mechanism by which SpCas9 and SaCas9 promote DNA melting and RNA-DNA heteroduplex formation—a process for which an atomic-level description has been incomplete. First, we observed in simulations that sgRNA-loaded Cas9 can bind and bend dsDNA in the groove between the REC and NUC lobes. Consistent with this observation, mutation of residues that mediated DNA binding in our simulations altered Cas9 editing efficiency in a cell-based gene editing assay. Notably, the interlobe cleft is not only a conserved feature within the Cas9 family, but is also a shared structural feature of the distantly related Cas12^40^ and Cascade^41^ enzymes. Binding of dsDNA in the interlobe space, where the DNA is adjacent to the sgRNA, could thus be a common feature of Cas-dsDNA encounter complexes formed across the entire Cas family of enzymes.

We unexpectedly observed in subsequent simulations that sgRNA can enter into Mg^2+^-mediated interactions with the DNA that lead to the intercalation of sgRNA bases into the DNA and promote local and reversible DNA base-pair breaks. Consistent with this result, we found experimental evidence that an ssRNA, in the absence of Cas9, can significantly lower the melting temperature of dsDNA. Whereas prior models of Cas9-mediated dsDNA melting have hypothesized that dsDNA base-pair breaks occur spontaneously when dsDNA adopts a bent structure, our data provide evidence for an alternative hypothesis that interactions with sgRNA induce dsDNA melting. If, as conjectured earlier, binding of dsDNA in the interlobe space is a conserved feature of encounter complexes across the Cas family, RNA-mediated DNA melting might also be a general mechanism of DNA deformation adopted by these enzymes.

In our MD simulations, we also observed unidirectional, sequential RNA-DNA hybrid formation mediated by Cas9. Although MD simulations have been used previously to study specific aspects of Cas9 function,^42–49^ the simulations presented here provide the first dynamic view of Cas9-mediated heteroduplex formation at the atomic level. In addition, we observed that open-and-closed breathing motions of the cleft between the REC and NUC lobes fine-tuned the distance between bases in the tsDNA and sgRNA prior to hybrid formation, promoting in-register RNA-DNA base pairing starting from the PAM +1 position, and breaking nonproductive base pairs if formed—an apparent proofreading mechanism to ensure sequential base pairing from the immediate PAM proximal base. (We note that although our results strongly suggest that heteroduplex formation is initiated at +1, they are not inconsistent with a scenario in which heteroduplex formation is occasionally initiated at other sites.) It seems possible that insights gained in this work could contribute to modifying the efficiency and specificity of Cas9-based gene editing tools. Because the closed-to-open transition is important for Cas9 to ensure that hybrid formation is initiated from the PAM +1 position, we speculate that mutations in Cas9 that alter the kinetics of the breathing motions may alter the ability of Cas9 to correct nonproductive structures formed by partially mismatched sequences.

The sgRNA-Cas9-dsDNA encounter complexes discussed in this work include an open-Cas9 conformation bound to a bent dsDNA that resembles the post-heteroduplex-formation structures obtained through X-ray crystallography (PDB ID 5F9R^17^ for SpCas9; PDB ID 5CZZ^29^ for SaCas9). Notably, similar encounter complex structures can be independently obtained using Boltz-1,^48^ an open-source deep-learning model for predicting structures of biomolecular complexes that has a similar architecture and accuracy to AlphaFold3 (see Supplementary Discussion and Fig. S5, A and B). Despite the high consistency between models predicted by two orthogonal computational modeling methods, our study does not rule out the possibility that other Cas9-dsDNA encounter complexes exist, and that Cas9-mediated DNA-RNA hybrid formation could happen by alternative pathways that may not be related to the mechanism discussed here.

In our PAM-restricted simulations, we observed that dsDNA can form encounter complexes with closed-SpCas9 conformations through interactions with the REC2 domain and PI domain, despite the fact that these complexes have fewer DNA-protein contacts than the observed open-Cas9–dsDNA complexes (Fig. S1D). A cryo-EM structure of SpCas9 covalently linked to a dsDNA^22^ (PDB ID 7S36) revealed that the protein can accommodate dsDNA binding in a similar closed conformation. In this structure, the dsDNA is captured in a severely bent and twisted conformation, with missing density in the PAM-proximal region of the DNA, consistent with the idea that a partial dsDNA base-pair break could happen immediately adjacent to the PAM motif and eventually progress to dsDNA-sgRNA hybrid formation. In simulations we initiated from this structure, the dsDNA was stable in the absence of the covalent linkage, although it remains to be established how productive R-loop formation could occur from this twisted and bent conformation (see Supplementary Discussion and Fig. S5). In our encounter simulations, dsDNA bound to closed-Cas9 remained straight, but this could represent an intermediate structural state prior to reaching the bent conformation seen in the cryo-EM structure. In general, it is likely that both local unwinding from the twisted and bent dsDNA conformation and RNA-mediated DNA melting contribute to dsDNA melting. Further experimental work with SpCas9 and other systems will be needed to determine the relative contributions of these mechanisms, which likely depends on the specific structural features of the enzyme in question.

A novel aspect of our proposed R-loop formation mechanism is the involvement of a molten-helical state of dsDNA, characterized by reversible, transiently stable strand-separated structures. Although direct experimental evidence for this state has yet to be established, our simulations highlight the potential significance of such structures in facilitating rapid flipping of tsDNA bases—a critical prerequisite for RNA-DNA heteroduplex formation. Base-flipping is promoted because partial melting reduces key energetic and structural barriers, such as base-stacking interactions and constraints on phosphate-backbone geometries, that typically stabilize the DNA double helix.

In summary, the results presented here provide strong evidence that an RNA-mediated, Mg^2+^-dependent mechanism contributes to DNA melting in structurally distinct Cas9 enzymes, and potentially other Cas enzymes. It will be interesting to determine whether such a mechanism, in which an ssRNA directly contributes to dsDNA melting, is important in non-Cas protein-nucleotide complexes, such as Argonaut-like systems.

## Acknowledgments

The authors thank Michael Eastwood for helpful discussions and Eric Martens for editorial assistance.

## Methods

### Structure preparation

The initial conformations in our simulations of SpCas9-sgRNA were built from a crystal structure of the pre-target protein-sgRNA complex (PDB ID 4ZT0^20^). The initial conformations of SaCas9-sgRNA were built from the post-hybrid complex crystal structure (PDB ID 5CZZ^29^). DNA was removed from the post-hybrid structures, and any co-crystallized ions (e.g., Mg^2+^, PO_4_^3−^, and SO ^2−^), water molecules, or buffering reagents were removed. Termini of the proteins were capped with ACE/NME capping groups. Termini of the RNA were not capped.

Missing protein loops were modelled based on underlying crystallographic density using Coot.^50^ Missing atoms were added using the Protein Preparation Wizard in Maestro (Schrödinger, LLC). Linear B-form DNA duplexes were built de novo in Coot, and the two DNA sequences complemented the corresponding sgRNA sequences in SpCas9 and SaCas9. Termini of the DNA were not capped. (See Table 1 for sequence information).

**Table 1.**
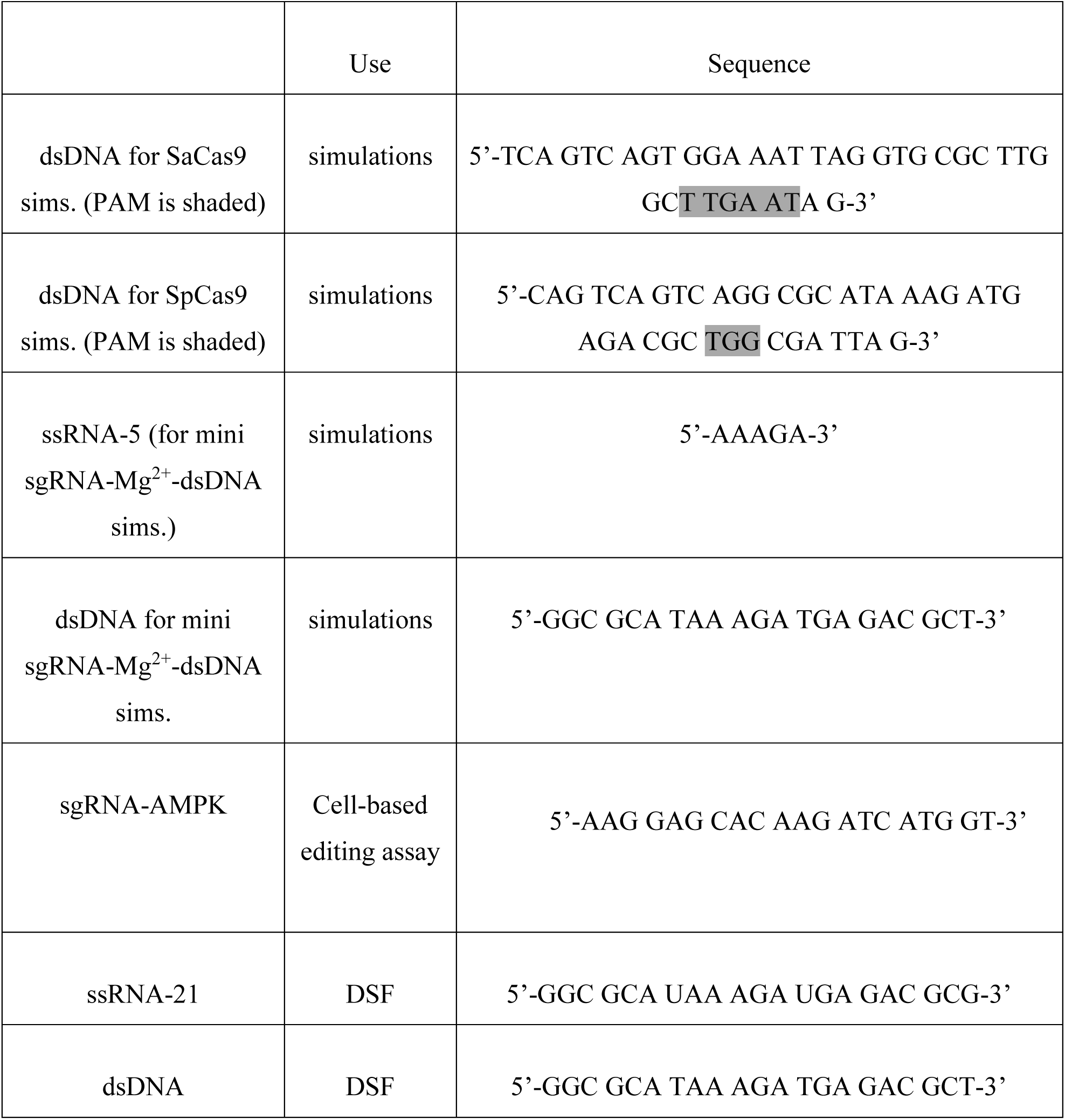
Nucleotide sequences.

### Parameterization

Proteins, ions, and nucleic acids were parameterized with the DES-Amber 3.20 force field.^51^ The force field was based on the previously published DES-Amber force field^52^ with changes to the phosphate group that improve the description of the interaction with proteins and result in more stable protein/nucleic acid complexes. The systems were solvated with water parameterized using the TIP4P-D water model,^53^ 0–25 mM MgCl_2_ and neutralized with a 150 mM NaCl. A typical system containing DNA-Cas9 contained ∼360,000 atoms in a 140 × 140 × 140 Å cubic box.

### Simulation details

Systems were first equilibrated on GPU Desmond^54^ using a mixed NVT/NPT schedule, followed by a 1 µs relaxation simulation on Anton, a special-purpose machine for conducting molecular dynamics simulations.^55^ Harmonic position restraints were applied to protein and nucleic acid backbone atoms during equilibration, and removed during the relaxation and production simulations. All production simulations were performed on Anton and initiated from the last frame of the relaxation simulation. Production simulations were performed in the NPT ensemble^56^ at 310 K and 1 atm using the Nosé-Hoover thermostat^57^ and the Martyna-Tobias-Klein barostat,^58^ respectively, applied using the multigrator framework.^59^ The simulation time step was 2.5 fs, and a modified r-RESPA integrator^60^ was used in which long-range electrostatic interactions were evaluated every three time steps. Electrostatic forces were calculated using the *u*-series method.^61^ A 9 Å cutoff was applied for the van der Waals calculations. The detailed setup and specific restraints used in simulations, if any, are specified below.

#### Free-binding simulations

For the free-binding simulations, the dsDNA was manually placed in the simulation box at one of two different locations, with a minimal distance between the dsDNA and Cas9-sgRNA complexes of more than 20 Å. For each dsDNA position, 10 replicate simulations (by which we mean simulations having the same starting coordinates but different thermally randomized starting velocities) were of 5 μs were performed.

#### PAM-restrained simulations

We conducted 20 10 μs PAM-restrained simulations (10 replicate simulations each for SaCas9 and SpCas9) for which the dsDNA was manually placed above the REC-NUC groove of Cas9, with the PAM motif within 5 Å of the PAM-binding arginines (Arg 1015 in SaCas9; Arg 1333 and Arg 1335 in SpCas9) in the PI domain. The rest of the DNA was more than 3.5 Å away from the protein-RNA complexes. The distance between the PAM-coordinating arginine and the corresponding DNA bases was restrained by a harmonic potential force (spring constant: 0.35 kcal mol^−1^ Å^−2^).

#### Simulations of dsDNA-Cas9 complexes

For each of SaCas9 and SpCas9, a dsDNA-Cas9 encounter complex in which the DNA was bound in the interlobe groove and bent was extracted from the PAM-restrained simulations, reparameterized, and used to initiate 20 replicate simulations of up to 50 μs. In this set of simulations, the distance between the PAM-coordinating arginine(s) and the corresponding DNA bases of the PAM was restrained by a harmonic potential force (spring constant: 0.35 kcal mol^−1^ Å^−2^), and the distance between each of three terminal base pairs at each end of the dsDNA were also restrained by a flat bottom well potential force (spring constant: 80 kcal mol^−1^ Å^−2^) to prevent the expected fraying motions due to the partially broken stacking interactions at the termini of the dsDNA (which does not happen in the cellular context, in which Cas9 binds in the middle of genomic DNA).

#### Simulations of isolated dsDNA-ssRNA complexes and dsDNA alone

The initial conformation of the mini ssRNA/dsDNA complex comprising ssRNA-5 and the 20 nt dsDNA was extracted from a simulation of a 40 nt dsDNA–sgRNA-SpCas9 complex in which the Mg^2+^-mediated dsDNA-sgRNA interaction and the RNA base minor groove intercalation was observed. The mini complex was resolvated, neutralized, and parameterized, and 20 replicate simulations of at least 20 μs and up to 100 μs were performed. For the simulations with dsDNA alone, the ssRNA was removed from the complex, and the 20 nt dsDNA was resolvated and reparameterized. Three replicate simulations of 100 μs were performed.

#### Simulations that temper the DNA interstrand interactions

Enhanced sampling methods have frequently been used to sample conformational states that can only be accessed on long timescales, and have been shown to be helpful in studying SpCas9 conformational plasticity.^62,63^ To enhance the sampling of DNA base-pair breaks in our Cas9-bound DNA simulations, we used Times Square sampling (TSS)^64^ to temper the strength of the interactions between the DNA strands. In our TSS simulations, which we also refer to as our tempering simulations, the strength of the near electrostatic interactions between the positively charged atoms on one DNA strand and the negatively charged atoms on the other DNA strand (and vice versa) were scaled by a factor λ. Λ was only allowed to adopt certain discrete values, sometimes referred to as “rungs,” including λ = 1 (the first rung, where the interactions are normal) as well as certain values λ < 1 (where the interactions are weakened) down to λ = λ_min_ at the final rung. The value of λ fluctuates over the course of the TSS simulation (tempering the interactions of the DNA strands), but the rules for changing rungs ensure that the structures sampled whenever λ = 1 are sampled from the same Boltzmann distribution as in a conventional MD simulation with normal interactions. (TSS is related to simulated tempering,^65,66^ but adaptively updates estimates of inter-rung free energy differences on-the-fly, allowing it to move more efficiently between rungs; its use in a related problem, tempering the interactions between proteins, has been described previously.^67,68^) We varied the tempering strength by choosing different values for λ_min_ from 0.980 to 0.995, and found that the sampling efficiency was not sensitive to λ_min_ in this range. Λ_min_ = 0.985 was used for the bulk of the SpCas9 simulations, and λ_min_ = 0.980 was used for the SaCas9 simulations.

The initial dsDNA-Cas9 encounter complexes were extracted from the PAM-restrained simulations. Scaling was applied only for the 20 DNA base pairs downstream of the PAM, starting from +1. DES-Amber 3.20 contains a charge scaling that makes it currently incompatible with charge tempering in Anton simulations, so for these simulations we used DES-Amber SF 1.0, which is similar but does not contain this scaling. Notably, the overall shape of the histograms of first base-pair break events under TSS resembles what we observed in general non-tempering simulations (Fig. S4A and Fig. 2D), suggesting that TSS preserves essential features of position-dependent DNA breaks. In addition, melting (as defined by multiple consecutive base-pair breaks in the DNA) was reversible in our TSS simulations, creating partially melted DNA structures that we termed the “molten-helical” state of DNA (Fig. 4 and Fig. S4, A and B).

### Simulation Analysis

#### Visualization

Simulation trajectories were visualized using a modified version of Visual Molecular Dynamics (VMD).^69^ For trajectory plots (e.g., for DNA bend angle, RNA-DNA distance, etc.), the raw measurement is shown in cyan, and the mean of a sliding window average of 10 data points is shown in red. Structure images were rendered using PyMOL (Schrödinger, LLC).

#### Protein-DNA contact analysis

Protein-DNA contact analysis was performed using a 3.5-Å cutoff to define a contact, and by the number of protein-DNA contacts we mean the number of protein residues in contact with the DNA. (If any number of atoms of a given amino acid residue came within the contact distance of at least one atom of any DNA base, then a contact was counted for that residue.) For the free-binding simulations, 1000 frames, evenly distributed within the simulation, were analyzed from each simulation.

#### dsDNA heteroduplex formation plot and melting plot

The intra–base pair distance at each position of the tsDNA-sgRNA hybrid (for heteroduplex formation) and the dsDNA (for melting) was defined as the distance between the heavy atoms of the NH-N hydrogen bond common to both A-U (A-T) and C-G base pairs.

### Cell-based gene editing assay

The cell-based gene editing assay was conducted by the contracted research organization HD Biosciences in San Diego. Mutations in FLAG-tagged SpCas9 were introduced in the pLentiCRISPR v2 (Addgene #52961)^70^ by point mutagenesis, and the mutated sequences were validated by sequencing. The vectors used in the study all encode an sgRNA complementary with the human AMPK B2 gene (see Table 1 for the sgRNA sequence), which has previously shown to be robustly edited using the protocol developed at HD Biosciences.

Human embryonic kidney (HEK) 293T cells were cultured in DMEM (ThermoFisher, Cat#11995-065) with 10% FBS (Omega Scientific Inc., Cat#FB-02) and 1% P/S (ThermoFisher, Cat#15140-122), and transfected with the WT and mutant CRISPR systems using a previously reported protocol.^71^ Transfected cells were selected by the inclusion of 1 µg/mL puromycin in the medium. A plasmid expressing GFP was co-transfected to monitor the transfection efficiency, which was measured by flow cytometry 24 h after transfection; a typical experiment had greater than 70% GFP-positive cells. Untransfected cells served as negative controls. Three days after transfection, cells were collected and each sample was split into two parts. Half of the cells were used to assess SpCas9 expression by western blot analysis using both an anti-Flag antibody (Biolegend, Cat#637301) and an anti-SpCas9 antibody (Biolegend, Cat#844302), and the results from the two antibodies were consistent. From the other half of the cells, genomic DNA was extracted for next-generation sequencing (NGS) to assay the on-target editing efficiencies of the WT and mutant proteins. The results were normalized using the western blot data, and results were averaged from experiments conducted on two different days.

### Differential scanning fluorimetry analysis

The ssRNA and dsDNA were synthesized by Ella Biotech. The DNA duplex was annealed and the product was substantially purified by HPLC. In order to identify melting transitions in the DNA, differential scanning fluorometry (DSF) analysis was conducted by the contract research organization Crelux in Munich, Germany. Briefly, fluorescence of SYBR Green I (Invitrogen) was measured on QuantStudio 7 Flex (ThermoFisher, Cat#4485701) in buffers containing 50 mM Tris-HCl pH 7.5 and 150 mM NaCl supplemented with different concentrations of MgCl_2_ and RNA/DNA (see corresponding text or figure for specific conditions). The samples were subjected to a temperature ramping from 27 °C to 95 °C at a rate of 2 °C/min. The derivatives of the raw fluorescence curves were used to identify the transition events, and the melting temperature of the dsDNA was determined by the peak temperatures. The data analysis was conducted according to the instructions provided by the device manufacturer.

### Structure predictions by Boltz-1

Boltz-1 (v0.4.1)^48,72^ predictions were performed locally on a commodity GPU cluster, with all parameters set to default. The protein and nucleic acid sequences used for each co-folding run are listed in Table 2. The multiple sequence alignments were generated using hhblits from HH-suite3,^73^ with all parameters set to default. The five highest ranking models predicted from each run were manually inspected (see Supplementary Discussion for details).

**Table 2.**
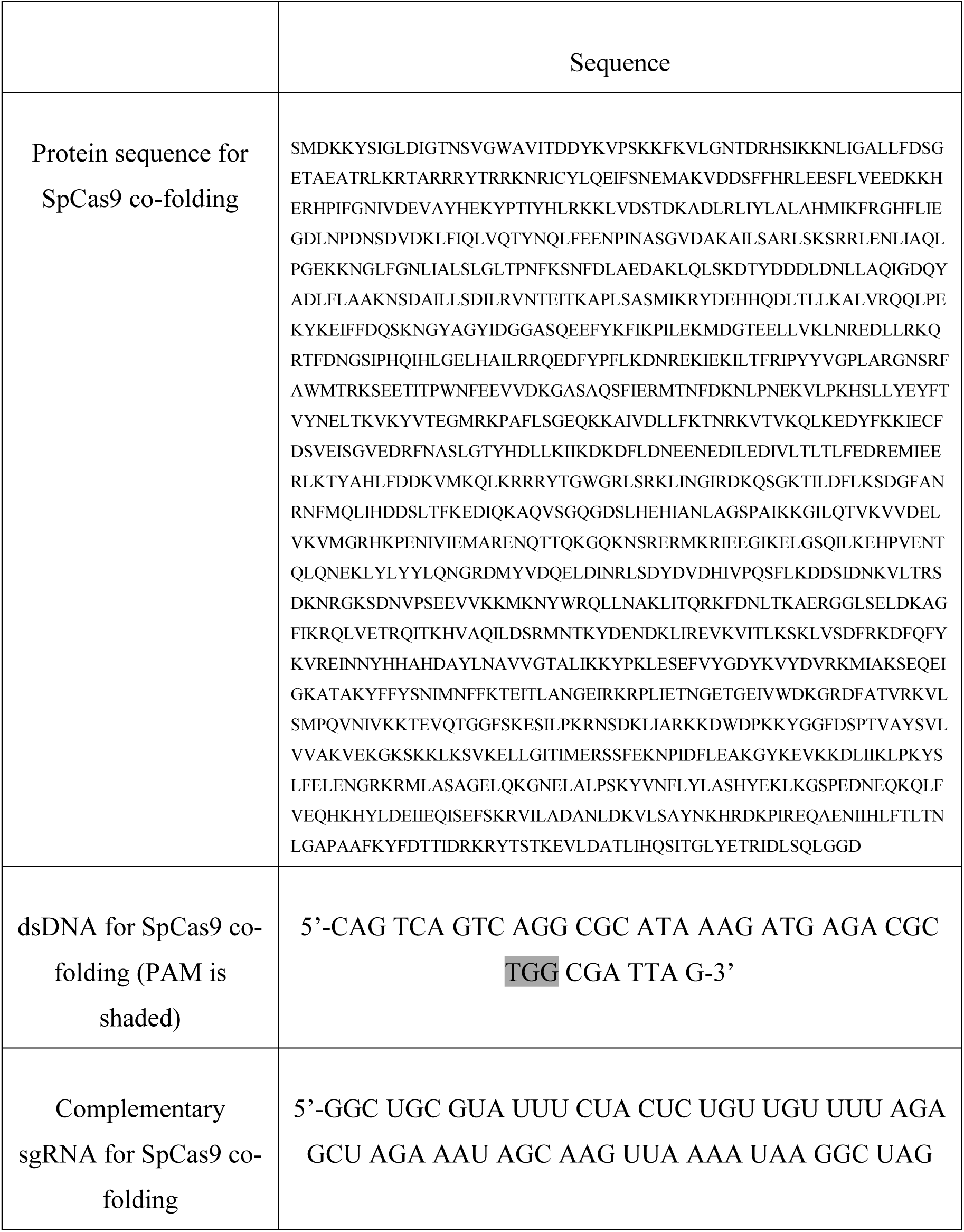
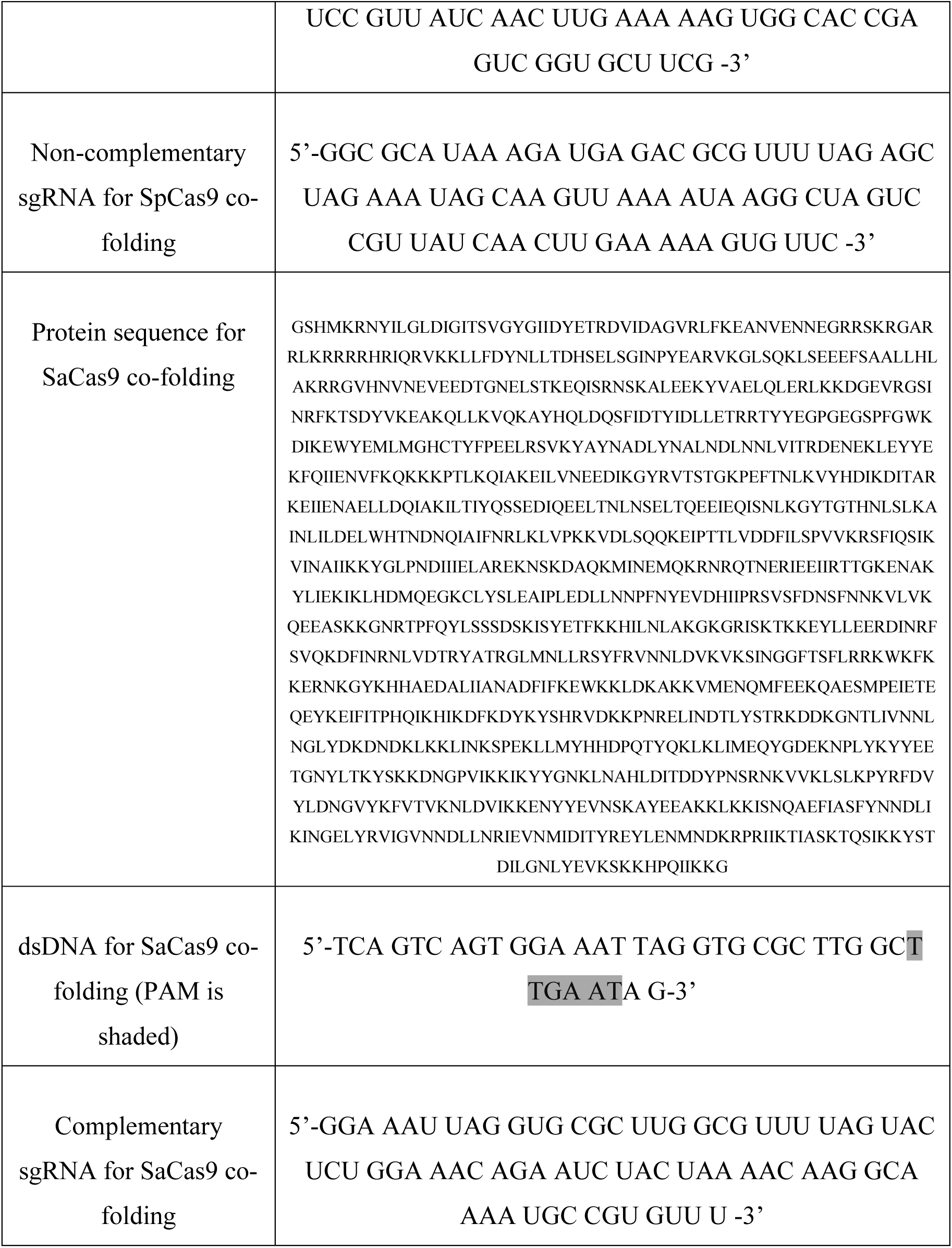
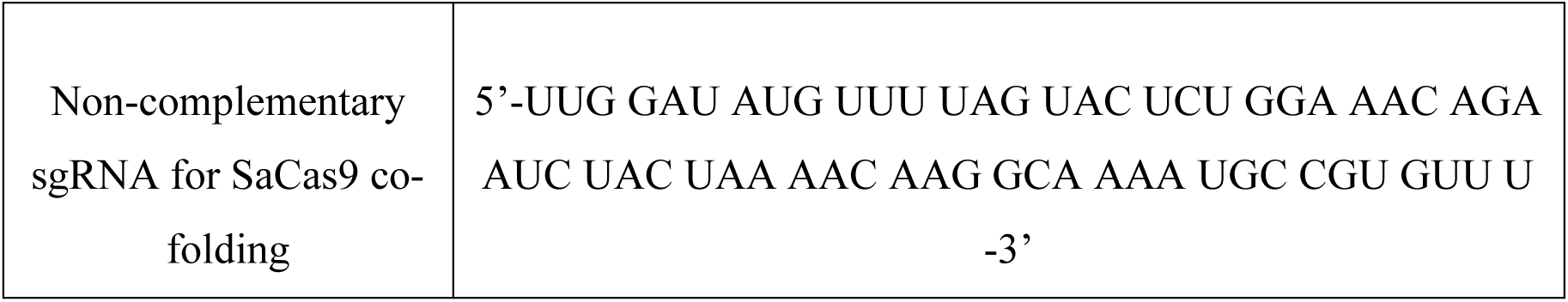
Protein and nucleic sequences for Boltz-1 structure predictions.

