## Supplementary Information for "An RNA-mediated DNA melting mechanism for CRISPR-Cas9"

### Supplementary Discussion

#### *dsDNA-sgRNA-Cas9 complex structure predictions by Boltz-1*

We first used Boltz-1 to co-fold three sequences: SpCas9, a 40-bp dsDNA containing a PAM motif, and a 20-base sgRNA with a sequence complementary to the region downstream of the PAM motif in the dsDNA (the sequences were the same as those used in our encounter simulations). The five highest-ranking models, which were highly similar, were manually inspected (we refer to them as the SpCas9-Boltz-A models), and the top-ranked model was used for RMSD comparisons with other structures. The overall conformation of the SpCas9-Boltz-A models closely resembles that of SpCas9 in the crystal structure of the post-hybrid state (PDB ID 5F9R), with a backbone Cα RMSD of 2.04 Å between the two structures. By comparison, alignment of the Boltz-1 models to the dsDNA-free closed-Cas9 state (PDB ID 4ZT0) results in an RMSD of 4.75 Å, and alignment to our MD-generated dsDNA-bound open-Cas9 state yields an RMSD of 4.79 Å (Fig. S5A). Specific similarities to the PDB 5F9R crystal structure include correct engagement of the PAM sequence with Arg1333 and Arg1335 of the PI domain, and formation of an RNA-DNA heteroduplex by the 17 tsDNA nucleotides downstream of the PAM motif. These structural features support the notion that the SpCas9-Boltz-A models adopt a post-hybrid, pre-cleavage state (Fig. S5A). It should be noted, however, that closer inspection of the models revealed numerous unrealistically close atomic contacts, suggesting that these models are not suitable for detailed analysis at the atomic level. Despite these limitations, the SpCas9-Boltz-A models successfully capture the essential features of the post-hybrid state of the dsDNA-sgRNA-SpCas9 complex.

Next, we replaced the guide sequence in the sgRNA with a randomly generated sequence that lacked complementarity to the dsDNA. The five highest-ranking models, which we refer to as the SpCas9-Boltz-B models, were again highly similar. We found that they closely resemble our

MD-generated dsDNA-bound open-Cas9 conformation, and alignment of the highest-ranking Boltz-1 model with our MD structure results in a backbone Cα RMSD of 3.49 Å. By comparison, alignment with the dsDNA-free closed-Cas9 conformation (PDB ID 4ZT0) results in an RMSD of 5.23 Å, and alignment with the post-hybrid state (PDB ID 5F9R) yields an RMSD of 4.49 Å (Fig. S5A). Upon closer examination of the SpCas9-Boltz-B models, we found that the sgRNA is unpaired, and the dsDNA is positioned in the groove between the REC and NUC lobes, adopting a bent conformation similar to that observed in our MD simulations. Notably, in these Boltz-1 models, the sgRNA backbone lies in close proximity to the dsDNA, particularly at the convex points PAM i+9 and PAM i+10—a feature that was likewise observed in our MD simulations.

The most significant difference between our DNA-bound, open-Cas9 models and the SpCas9-Boltz-B models is the location of the HNH domain. In the Boltz-1 models, the HNH domain adopts a position similar to that in the catalytically active enzyme, whereas in our MD-generated models, it is located in the position observed in the dsDNA-free closed-Cas9 conformation. This finding suggests that although the SpCas9-Boltz-B models represent a pre-hybrid state, it likely corresponds to a state adopted later in the encounter process than we sampled in our MD simulations.

Despite differences between our simulation models and the SpCas9-Boltz-B models, the latter capture three key features observed in our MD-generated complexes: (1) dsDNA bound deep within the groove between the REC and NUC lobes, (2) a bent dsDNA conformation with a convex point near PAM i+11, and (3) close contact between the dsDNA and sgRNA backbone. Using a similar protocol, we demonstrated that Boltz-1 predictions for SaCas9 yield comparable conclusions (Fig. S5B).

***Simulation analysis of bent and twisted dsDNA conformations bound to Cas9***

To probe the functional significance of the twisted dsDNA-Cas9 conformations observed in the cryo-EM study discussed in the main text (Ref. 22), we first performed 10 50-μs simulations initiated from one of these structures (PDB ID 7S36), with the covalent linkage between the DNA and protein removed. In all 10 simulations, we observed that at least one of the flipped tsDNA bases (PAM +1 or +2) reverted to Watson–Crick base pairing with its complementary DNA base, indicating that these flipped structures are only transiently stable (Fig. S5C). Although reversible transient breaks were also observed after Watson–Crick pairing was re-established, we did not observe the initiation of heteroduplex formation at any point during the 500 μs of total simulation time. We next performed five additional 10-μs simulations using a closely related cryo-EM structure (PDB ID 7S38), in which a nascent R-loop with three RNA-DNA base pairs had already formed, and the DNA was still in the twisted conformation. In all five simulations, at least one of the three RNA-DNA base pairs experienced frequent disruptions, and in one simulation, all three base pairs were completely broken by the end of the 10-μs trajectory (Fig. S5D). These results suggest that this early R-loop structure is inherently unstable under the simulation conditions, and it remains to be established how the R-loop could extend from this twisted dsDNA conformation. We speculate that additional large-scale protein conformation change may be needed to stabilize the heteroduplex, but this does not happen on the timescale of these simulations.

### Supplementary Figures

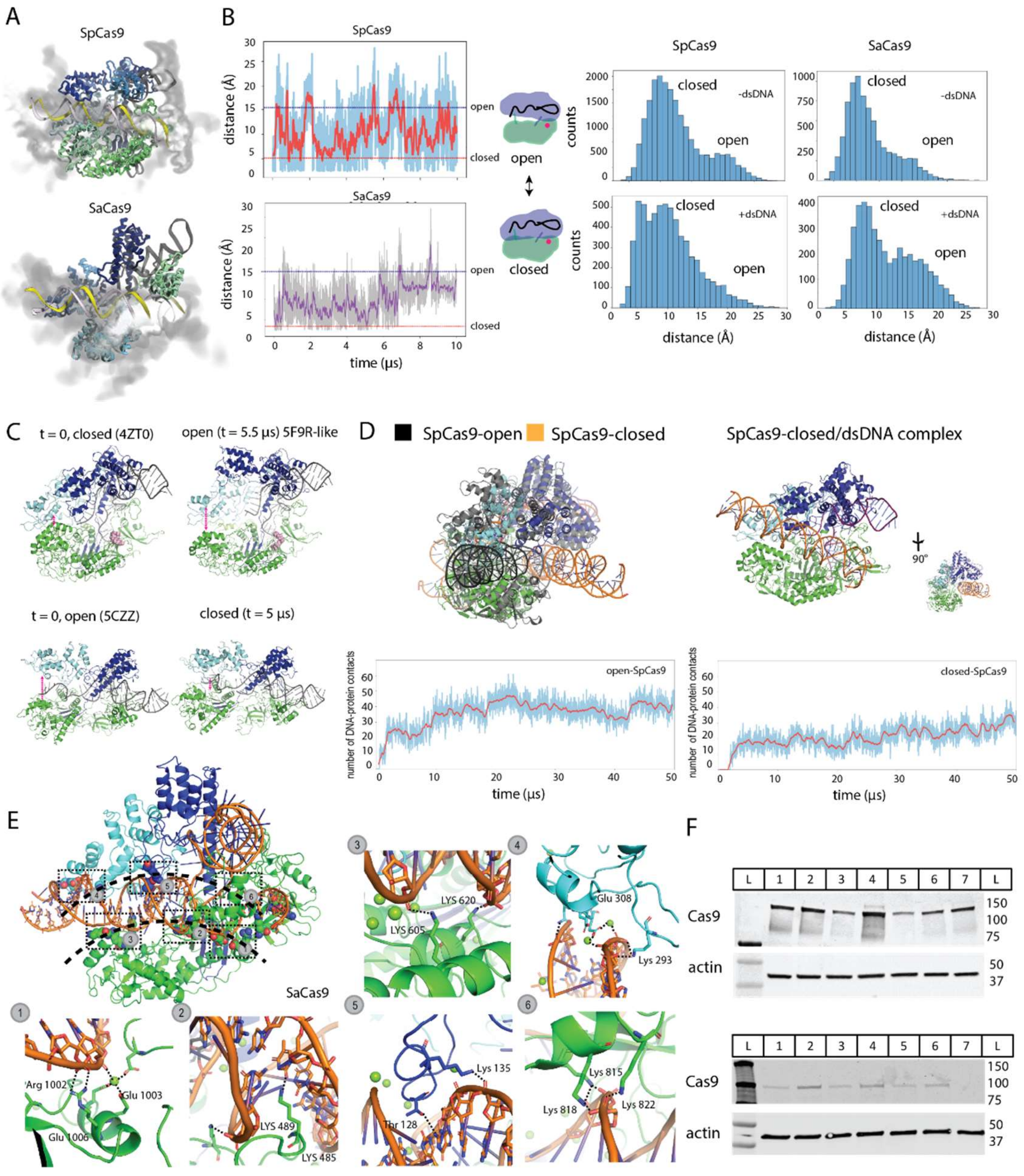


**Fig. S1. Simulation analysis of dsDNA-bound Cas9 complexes.** (A) Volume density maps showing the positions of bound dsDNA generated from all of the encounter complexes obtained from the free-binding simulations. The dsDNA in yellow and white indicates the pose in which the most DNA-protein contacts were formed. (B) Simulations of SpCas9 and SaCas9, in the absence of DNA, indicating reversible conformational transitions between their open and closed conformations. The plots on the left show the distance between the REC3 and RuvC domains during the course of the simulations. To the right of these plots is a simplified schematic of the open-and-closed motions of Cas9. On the far right are histograms of the distances between the REC3 and RuvC domains in sgRNA-Cas9-only simulations (top) and in sgRNA-Cas9-dsDNA encounter complexes generated from free-binding simulations (bottom). (C) Representative structural models of the open and closed Cas9 conformations. (D) Top: structural comparison of dsDNA bound to the open conformation of SpCas9 (DNA in black) and bound to the closed conformation of SpCas9 (DNA in orange). Bottom: PAM-restricted simulations in which dsDNA was bound to either an open (left) or closed (right) conformation of SpCas9 at the end of the simulation. The maximum number of contacts formed with the open SpCas9 conformation is higher than that with the closed conformation of SpCas9. (E) Atomic-level detail of how certain clusters of residues in SaCas9 interact with bent dsDNA. The black dashed lines indicate the concave surface of the NUC lobe. (F) Upper: Expression levels of WT SpCas9 and variants in HEK293 cells. L: molecular weight marker; 1: WT; 2: K233A/K234A; 3: R1333A; 4: K528A/K535A/K536A; 5: R778A/R780A/R783A/K782A; 6: K1200A/S1216A/Q1221A; 7: K1014A/Y1016A/R1019A/K1020A. Lower: Expression levels of WT SpCas9 and variants in HEK293 cells. L: molecular weight marker; 1: WT; 2: R1333A; 3: E779R/M781R/K782R; 4: K1014R/Y1016R/K1020R; 5: K233R/K234R/E232R/N235R; 6: K528R/K535R/K536R; 7: no transfection.

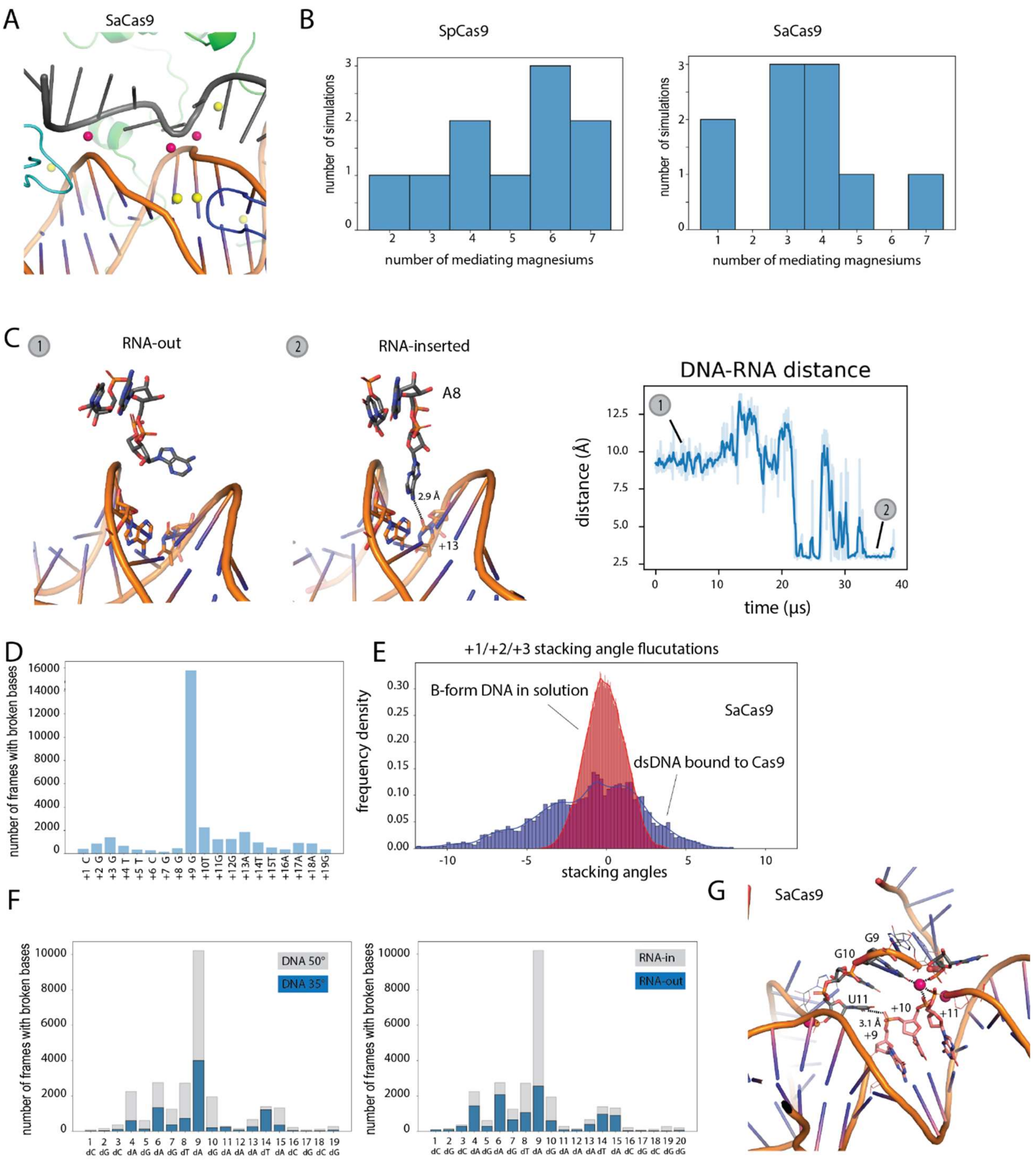


**Fig. S2. $Mg^{2+}$-mediated sgRNA-dsDNA interactions in dsDNA-Cas9 encounter complexes.** (A) A representative structure of $Mg^{2+}$ ions binding between the sgRNA (grey) and dsDNA (orange) backbones in an SaCas9 encounter complex. Pink spheres are $Mg^{2+}$ ions mediating backbone interactions; yellow spheres are $Mg^{2+}$ ions that do not mediate backbone interactions.

(B) Histogram distribution of the number of interaction-mediating $Mg^{2+}$ ions across Cas9-dsDNA encounter complex simulations.  (C) Left: Atomic-level representations of (1) a flipped RNA base and (2) the RNA base interacting with a dsDNA base in the minor groove in an SaCas9 encounter complex simulation.  Right: Simulation showing the minimal interatomic distance between A8 of the RNA and T(PAM +13) of the tsDNA.  (D) Distribution of base pair breaks in simulations along the SaCas9-bound region of the DNA.  (E) Distribution of stacking angles at the PAM-proximal region (+1, +2, +3) in simulations of SaCas9-bound DNA (blue) and DNA in solution (red).  (F) Distribution of base-pair breaks along the SpCas9-bound region of dsDNA under different simulation conditions.  Left panel: RNA inserted into the minor groove of a dsDNA with a 50° bend angle (grey) or a 35° bend angle (blue).  Right panel: a dsDNA with a 50° bend angle with RNA inserted (grey) or without RNA inserted (blue).  (G) A representative structure from the SaCas9 encounter complex simulations showing that RNA bases can interact with the phosphate backbone of the dsDNA.

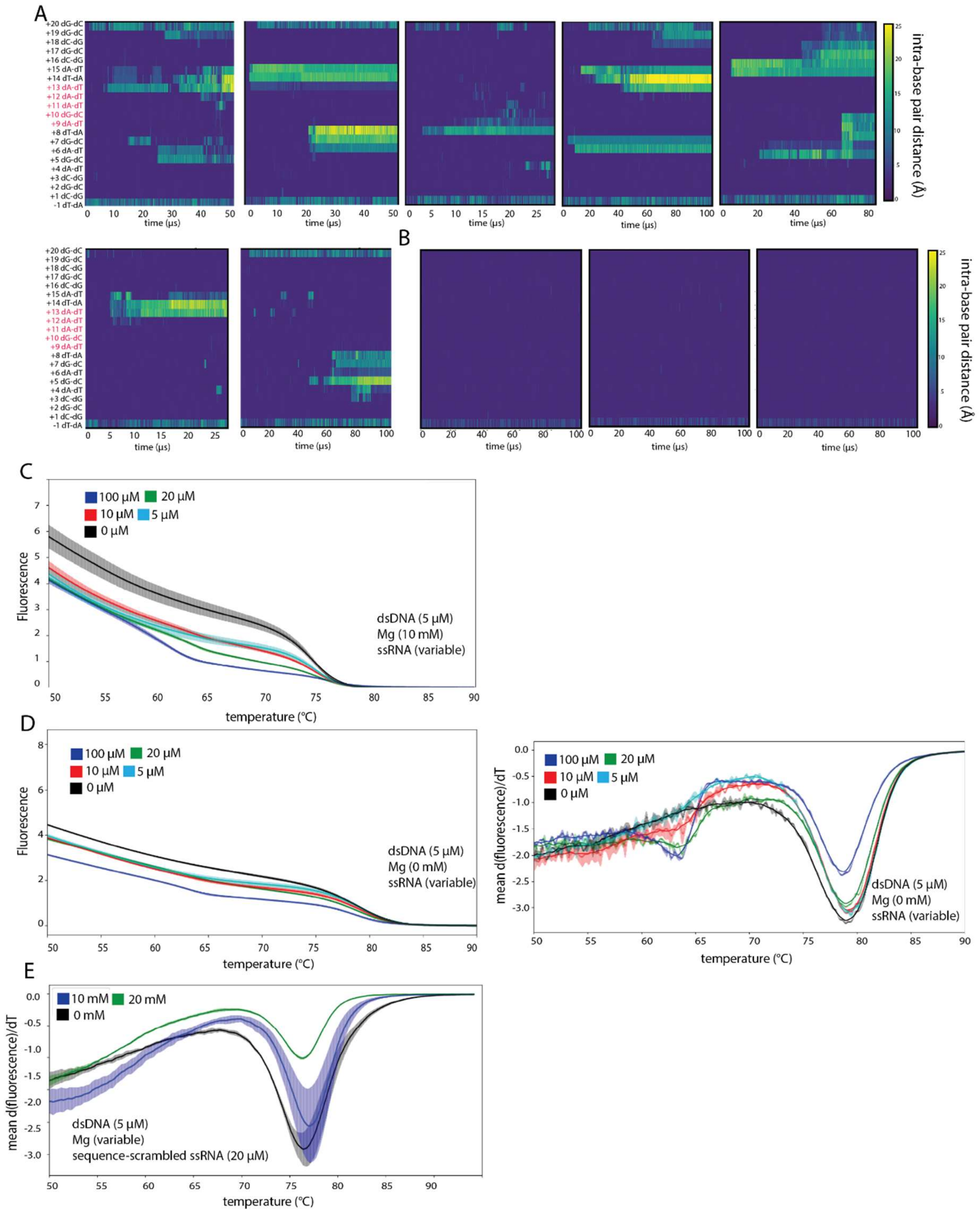


**Fig. S3. ssRNA promotes dsDNA melting.** (A) Base pair distance analyses, as in Fig. 3A bottom panel, but of seven additional simulations of dsDNA-ssRNA complexes. (B) Base pair distance analyses, as in Fig. 3A top right panel, but of three additional simulations of dsDNA in aqueous solution. (C) Raw fluorescence signal of dsDNA melting in the presence of 10 mM $Mg^{2+}$ and variable ssRNA concentrations. (D) Left: Raw fluorescence signal of dsDNA melting at variable ssRNA concentrations in the absence of $Mg^{2+}$. Right: Derivative of values plotted in the left panel. Standard deviations for the thermostability assays, which were performed in duplicate, are indicated by the lighter-colored bars around the mean. (E) Derivative of raw fluorescence signal of dsDNA melting in the presence of 20 μM sequence-scrambled ssRNA at variable $Mg^{2+}$ concentrations.

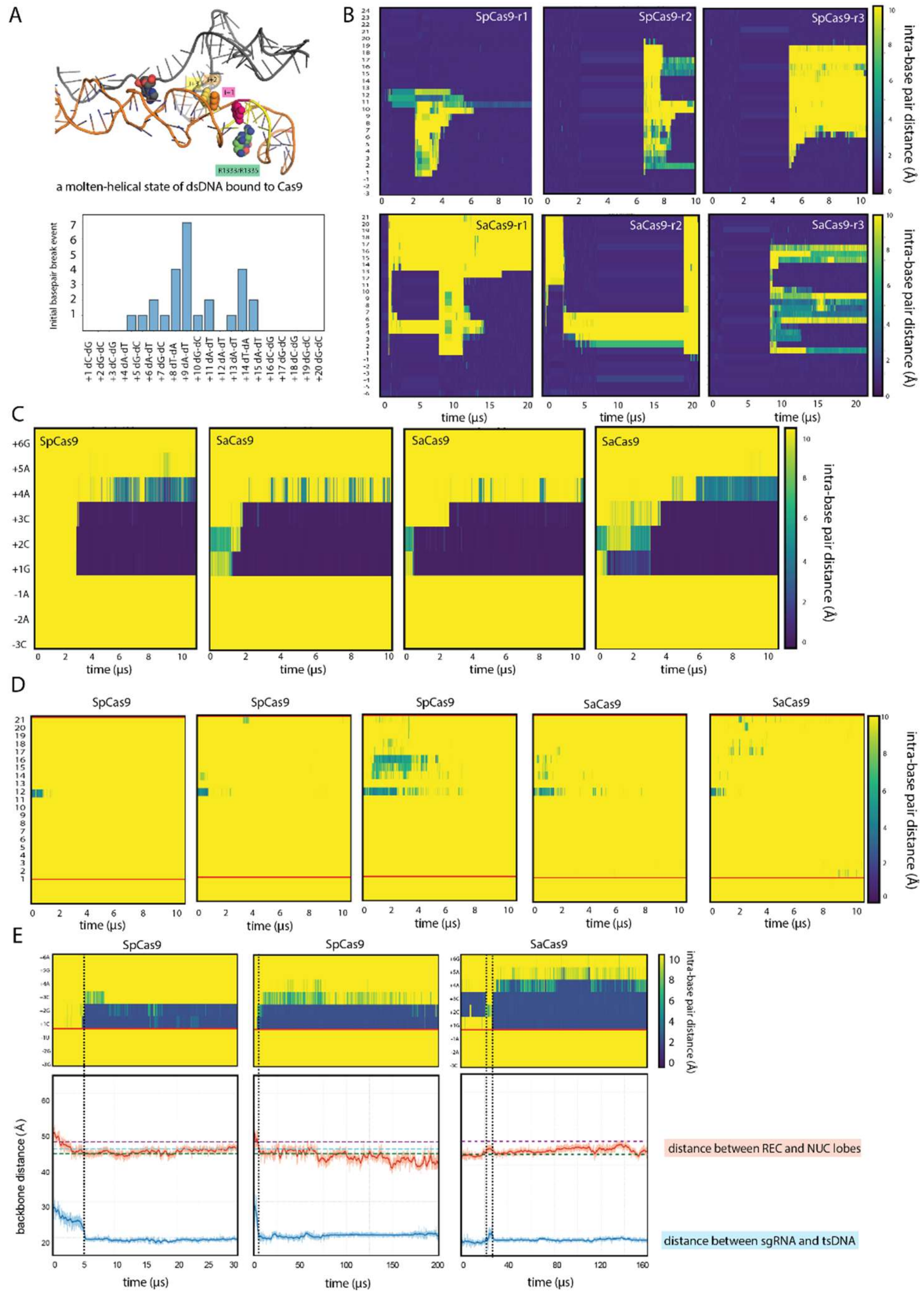


**Fig. S4. Heteroduplex formation in simulations of Cas9-dsDNA encounter complexes.** (A) Top: representative structure of a partially melted dsDNA bound to Cas9 in which the PAM-proximal bases have flipped out. The structure is taken from a simulation in which the electrostatic interactions between the two strands of the dsDNA were tempered. For Cas9, only the PAM-binding arginines are shown. Bottom: histogram of the first base pair to break in the tempering simulations. Although the tempering simulations allow dsDNA base pair breaks to be observed more rapidly, the distribution resembles what we observed with non-tempered simulation conditions, showing the use of tempering doesn't change which positions on the dsDNA are more likely to break first. (B) Base pair break analysis of three tempering simulations showing that reversible strand separation can be observed. Only frames from the first rung were used in this analysis in order to show that the reversibly formed molten-helical state was not the result of modifying the Hamiltonian. (C) Base pair distance analysis for each position in the tsDNA and the corresponding base on the sgRNA, measured as the distance between the heavy atoms of the NH-N hydrogen bond common to both A-T/U and C-G base pairs. Three SaCas9 simulations and one SpCas9 simulation are shown from the seven simulations total in which PAM +1–initiated heteroduplex formation occurred. (D) Base pair distance analysis for three simulations with SpCas9 and two simulations for SaCas9 in which the first heteroduplex pairing event happened in the PAM-distal region. The red lines indicate the position of PAM +1. (E) The bottom plots show backbone distance measurements between the center-of-mass of the phosphorous atoms of sgRNA bases +1, +2, and +3 (bound to the REC domain) and either the center of mass of the Cα atoms of the two PAM-binding arginines (in the NUC domain; red) or the center of mass of the PAM-proximal tsDNA bases (positions +1, +2, and +3; blue). Changes in these measurements reflect open and closed transitions of the REC and NUC domains. Motions that are correlated with changes in the corresponding sgRNA-DNA base pair distance analysis (top plots) are denoted by vertical dotted lines. The corresponding arginine-sgRNA distance from crystal and cryo-EM structures of SpCas9 are indicated as colored horizontal dotted lines (green: PDB ID 4ZT0; magenta: PDB ID 5F9R; cyan: PDB ID 6O0Z).

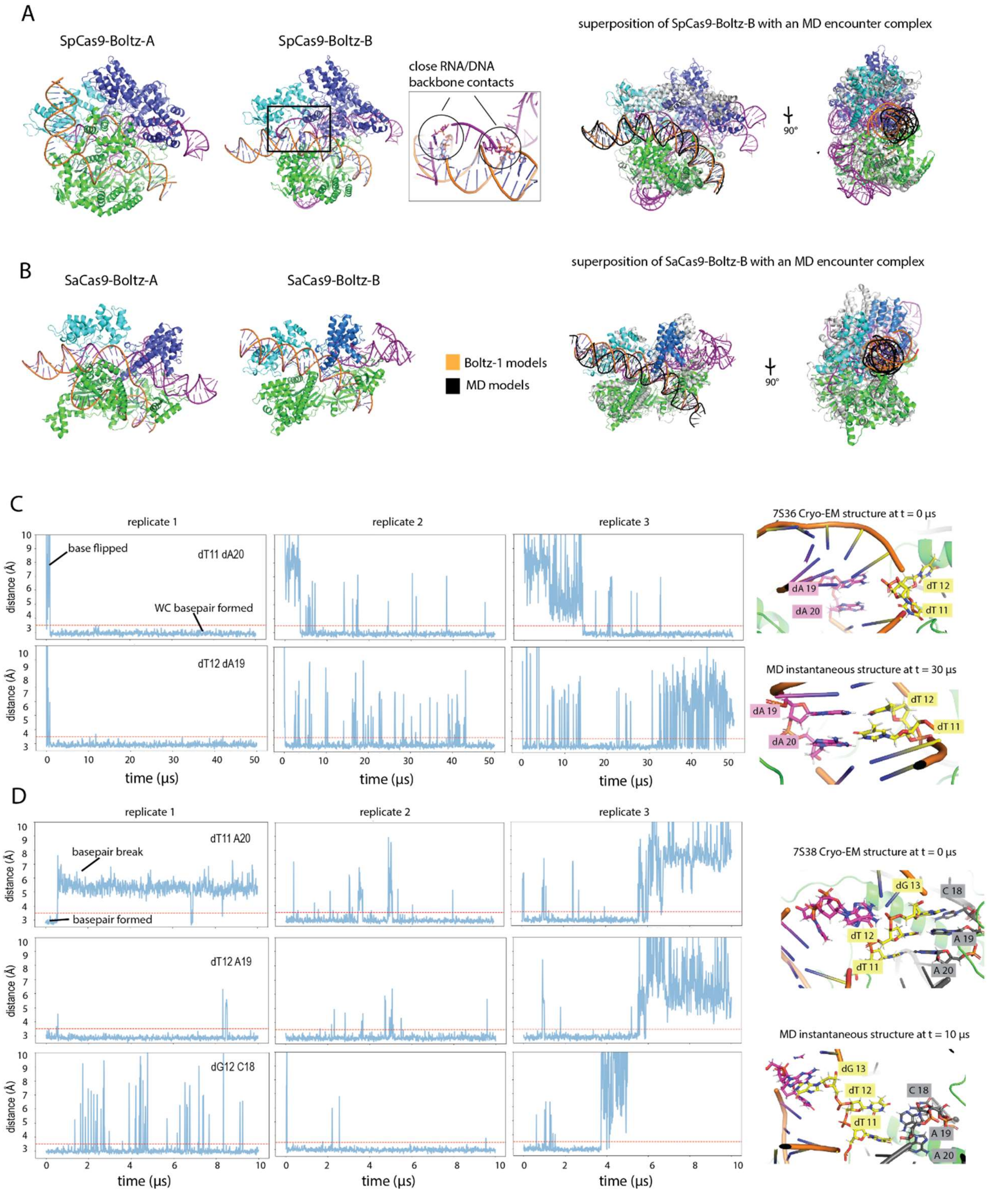


**Fig. S5. Boltz-1–predicted dsDNA-Cas9 encounter complexes and simulations of SpCas9-dsDNA cryo-EM structures with bent and twisted DNA conformations.** (A) dsDNA-SpCas9

encounter complexes with complementary sgRNA (SpCas9-Boltz-A) or noncomplementary sgRNA (SpCas9-Boltz-B) predicted by Boltz-1. The inset shows the close contact between the RNA and DNA in SpCas9-Boltz-B. On the right is SpCas9-Boltz-B superposed with a snapshot from a dsDNA-SpCas9 encounter simulation. (B) Left: dsDNA-SaCas9 encounter complexes with complementary guide RNA (SaCas9-Boltz-A) and noncomplementary guide RNA (SaCas9-Boltz-B) predicted by Boltz-1. Right: SaCas9-Boltz-B superposed with a snapshot from a dsDNA-SaCas9 encounter simulation. (C) Simulations of the SpCas9 cryo-EM structure with a bent and twisted dsDNA conformation prior to R-loop formation (PDB ID 7S36). Representative simulations showing the minimal interatomic distance between dT11(PAM +1) of the tsDNA and dA20 of the complementary DNA strand (upper), and the minimal interatomic distance between dA19 of the ntDNA and dT12(PAM +2) of the tsDNA (lower); Right: From one of these simulations, atomic-level representations of the starting structure and an instantaneous structure (at 30 μs) in which both dT11 and dT12 of the tsDNA formed Watson–Crick base pairs with the complementary DNA strand. (D) Simulations of the SpCas9 cryo-EM structure with a bent and twisted dsDNA conformation with a nascent three-base-pair R-loop (PDB ID 7S38). Representative simulations showing the minimal interatomic distance between A20 of the RNA and dT11(PAM +1) of the tsDNA (upper), the minimal interatomic distances between A19 of the RNA and dT12(PAM +2) of the tsDNA (middle), and the minimal interatomic distance between C18 of the RNA and dG13(PAM +3) of the tsDNA (bottom); Right: From one of these simulations, atomic-level representations of the starting structure and an instantaneous structure (at 10 μs) in which all three highlighted RNA-DNA base pairs are broken.

### Supplementary Video Captions

**Supplementary Video 1.** Formation of a dsDNA-SpCas9-RNA encounter complex in a free-binding simulation. DNA shown in orange, red, and blue; RNA shown in grey; SpCas9 shown in blue and green.

**Supplementary Video 2.** Straight-to-bent conformational change of dsDNA in a dsDNA-SpCas9-sgRNA encounter complex. DNA shown in yellow and white; RNA shown in light orange; Cas9 shown in blue and green. (The distance between the PAM nucleotides and the PAM-coordinating arginines was restrained during the simulations.)

**Supplementary Video 3.** $Mg^{2+}$-mediated sgRNA-dsDNA interactions observed in a simulation of a dsDNA-SpCas9-sgRNA encounter complex. DNA shown in yellow and white; RNA shown in light orange; Cas9 shown in blue and green; $Mg^{2+}$ shown as magenta spheres; intercalating RNA base depicted as a stick model. (The distance between the PAM nucleotides and the PAM-coordinating arginines was restrained during the simulations.)

**Supplementary Video 4.** ssRNA-5 facilitates melting in a 20-bp dsDNA. DNA backbone shown in yellow and white; RNA shown in light orange; $Mg^{2+}$ shown as magenta spheres; nucleic acid bases depicted as stick models.

**Supplementary Videos 5 and 6.** PAM +1–initiated, consecutive, and in-register sgRNA-tsDNA heteroduplex formation in a tempering simulation of a dsDNA-Cas9-sgRNA encounter complex (Video 5: SpCas9; Video 6: SaCas9). DNA backbone shown in yellow and white; RNA shown in light orange; nucleic acid bases depicted as stick models. +1 bases (of tsDNA and RNA) colored in magenta, +2 bases colored in orange, +3 bases colored in blue, +4 bases (Video 6 only) colored in green; coloring of bases changes from transparent to opaque when a base pair is formed. (In the tempering simulations the charged interactions between the two strands of the dsDNA were intermittently weakened, thereby increasing the frequency of DNA base pair breaks and accelerating the exploration of different DNA-DNA interactions.)

**Supplementary Video 7.** A tempering simulation of an SaCas9-dsDNA complex in which the base pairs first form out of sequence (i.e., at PAM +2 rather than +1) before the complex breaks the out-of-sequence base pairs and subsequently forms sequential base pairs from +1 to +4. Same color scheme as in Supplementary Video 6.
